# The molecular basis of tricalbin-mediated membrane contact site organization in cells

**DOI:** 10.1101/2025.07.11.664157

**Authors:** Lazar Ivanović, Anne-Laure Boinet, Andrea Picco, Marko Kaksonen, Wanda Kukulski

## Abstract

Membrane contact sites facilitate molecular exchanges through physical interactions between organelles, connected by specific protein tethers. Among these tethers are the tricalbins, which mediate contacts between endoplasmic reticulum (ER) and plasma membrane in yeast. Tricalbins are integral to the ER, have a cytosolic lipid binding domain and bind the plasma membrane through C2 domains. Here, we combine fluorescence recovery after photobleaching with correlative light and 3D electron microscopy to dissect how tricalbins control their localization, dynamic distribution and contact site organization. We find that heteromerization via lipid binding domains is a prerequisite for tricalbin accumulation at contact sites, membrane curvature sensing and restrained mobility in the ER. By altering tricalbin protein domains, we show that intermembrane distances and intrinsically disordered regions interdependently control distribution and dynamics of contact site tethers. Our study reveals principles of contact site architecture that are fine-tuned by tricalbin domain organization.

## Introduction

The endoplasmic reticulum (ER) interacts with other organelles in the cell via subdomains known as membrane contact sites, which are important for signalling, exchange of lipids, and organelle dynamics (Voeltz et al., 2024). In budding yeast, the contact sites between the ER and the plasma membrane, also referred to as the cortical ER (cER), appose approximately one third of the plasma membrane surface (Loewen et al., 2007; Manford et al., 2012; West et al., 2011). The connection between the ER and the plasma membrane is maintained predominantly by six conserved proteins from three families; two VAMP-associated proteins (VAPs) Scs2 and Scs22, TMEM16-like Ist2, and three tricalbins Tcb1, Tcb2 and Tcb3, orthologues of mammalian extended synaptotagmins (Manford et al., 2012).

Tricalbins preferentially associate with high curvature cER, dictated by a hydrophobic stretch that contains their ER transmembrane domain (Hoffmann et al., 2019) and bears structural similarity to ER-shaping reticulons (Giordano et al., 2013; Zurek et al., 2011) (Fig. 1A). Tricalbins also harbour a synaptotagmin-like mitochondrial-lipid-binding protein (SMP) domain which is positioned in the cytosolic gap and can transfer lipids between membranes (Bian et al., 2018; Giordano et al., 2013; Qian et al., 2021; Saheki et al., 2016; Schauder et al., 2014; Toulmay and Prinz, 2012). The SMP domain is followed by a sequence of up to six C2 domains which interact with anionic lipids in the plasma membrane (Jimenez and Davletov, 2007; Schulz and Creutz, 2004).

**Figure 1:**
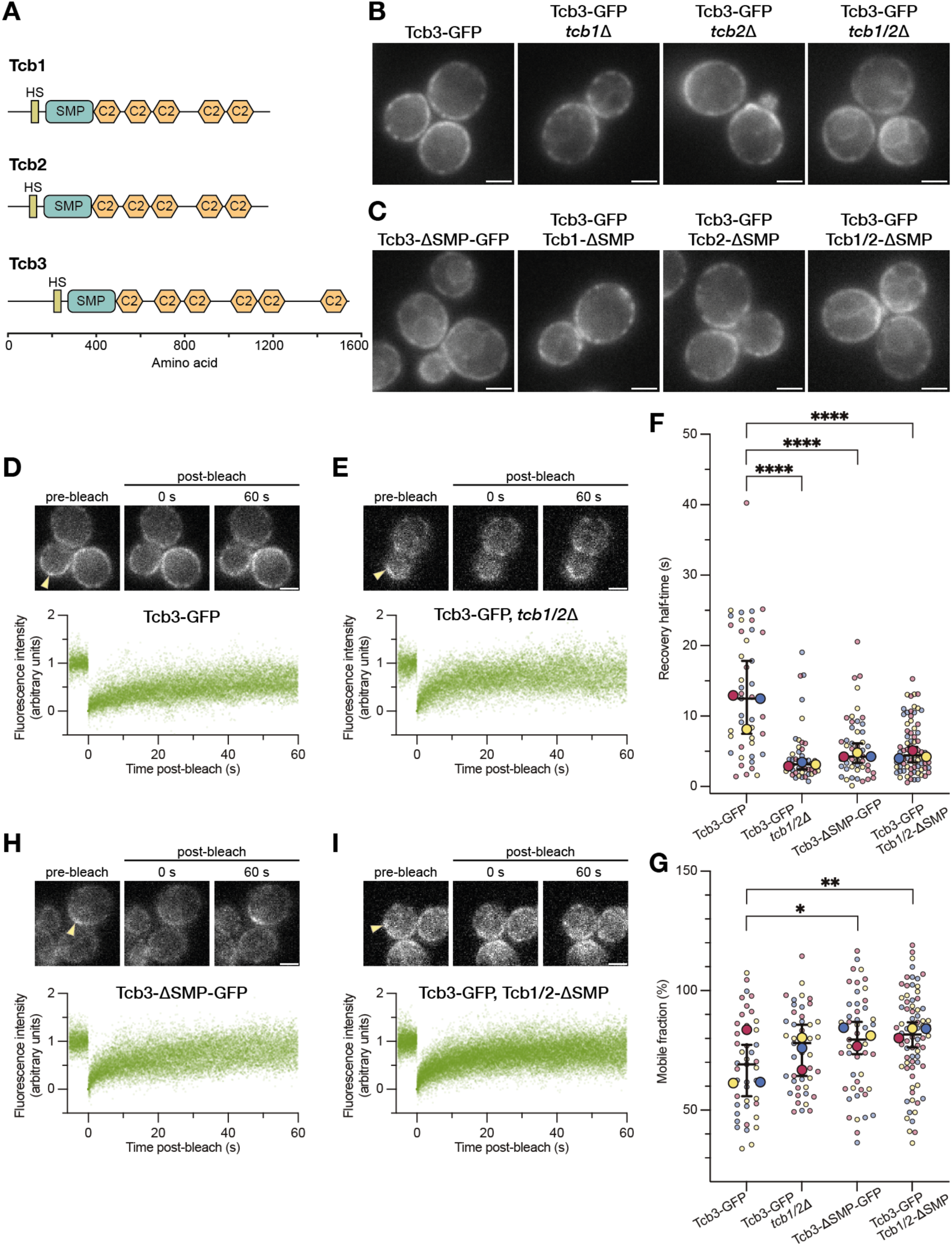
Cortical association and mobility of tricalbins depend on heteromerization via SMP domains. **A:** Domain organization of the tricalbins Tcb1, Tcb2 and Tcb3. HS: Hydrophobic stretch, which includes the transmembrane domain. C2 domains are based on AlphaFold2 predictions (Jumper et al., 2021). **B:** Fluorescence live cell imaging of Tcb3-GFP in WT, *tcb1Δ*, *tcb2Δ* and *tcb1/2Δ* cells. **C:** Fluorescence live cell imaging of Tcb3-ΔSMP-GFP in WT cells, Tcb3-GFP in combination with either Tcb1-ΔSMP, Tcb1-ΔSMP, or both Tcb1-ΔSMP and Tcb1-ΔSMP simultaneously. **D, E, H and I:** FRAP experiments on GFP-tagged Tcb3 variants in different genetic backgrounds, as indicated. The upper panels show images from representative FRAP movies, corresponding to time points before, at 0 s and 60 s after the bleaching event. Bleached regions are indicated by arrowheads. The lower panels show normalized and aligned fluorescence intensity measurements of all FRAP events (n=42 in D, n=43 in E, n=49 in H, n=76 in I, from 3 merged experimental repeats). **D:** FRAP of Tcb3-GFP in WT cells. **E:** FRAP of Tcb3-GFP in *tcb1/2Δ* cells. **F:** Recovery half-times and **G:** Mobile fractions determined from the FRAP experiments shown in D, E, H and L. The colors encode three experimental replicates. Each data point corresponds to an analyzed FRAP event. The larger points indicate the medians of each experimental repeat. The horizontal lines represent medians with 95% confidence interval of all data points combined. **H:** FRAP of Tcb3-ΔSMP-GFP in WT cells. **I:** FRAP of Tcb3-GFP in Tcb1/2-ΔSMP cells. Scale bars: 2 µm.

While the domain architecture of tricalbins explains their topology between the two membranes, it remains unclear how it contributes to retention and distribution of tricalbins within contact sites. Furthermore, the contact site ultrastructure associated with tricalbins is similar to that of VAPs and Ist2, and to a large extent the different cER proteins colocalize, despite being diverse in function and domain structure (Collado et al., 2019; Hoffmann et al., 2019). Thus, how the three tricalbins organize to mediate tethering, whether tethering modulates the intermembrane space between ER and plasma membrane, and how in turn the intermembrane space influences the behavior and organization of tricalbins, remains elusive. Here, we combine live cell imaging, fluorescence recovery after photobleaching (FRAP) and correlative light and electron microscopy (CLEM) to dissect molecular and architectural determinants of tricalbin-mediated membrane contact sites.

## Results

### SMP domain heteromerization drives tricalbin accumulation at ER-plasma membrane contact sites

To study the basis of tricalbin localization, we performed live cell imaging of GFP-tagged tricalbins. In wild type (WT) cells, Tcb3-GFP localizes to the cER (Manford et al., 2012; Toulmay and Prinz, 2012). We found that in absence of either Tcb1 or Tcb2, Tcb3-GFP maintained its exclusively cortical localization (Fig. 1B). Upon simultaneous deletion of both Tcb1 and Tcb2 (*tcb1**/**2*Δ), Tcb3-GFP was no longer exclusively cortical, but also appeared in internal ER, including the nuclear envelope (Fig. 1B)(Hoffmann et al., 2019). Previous work had shown that the SMP domains target tricalbins to the cER (Toulmay and Prinz, 2012). SMP domains are prone to dimerization (Jeong et al., 2017; Jeong et al., 2016; Kawano et al., 2018; Lees et al., 2017; Schauder et al., 2014), and tricalbins as well as extended synaptotagmins have been suggested to homo- and heteromerize (Creutz et al., 2004; Giordano et al., 2013; Hanaoka et al., 2024). Alphafold3 (Abramson et al., 2024) models of tricalbin pairs suggest dimeric SMP assemblies that are plausible given the arrangement of the SMP domains with respect to the predicted transmembrane regions (Fig. S1A). We therefore asked if the tricalbin SMP domains influence each other’s localization, which would indicate SMP heteromerization. We generated tricalbin SMP domain deletion mutants by CRISPR genome editing and live-imaged them. Tcb3-ΔSMP-GFP (residues 273-485 deleted) was distributed generically in all ER, similarly to full-length Tcb3-GFP in *tcb1**/**2*Δ cells (Fig. 1C). The same was true for Tcb1-ΔSMP-GFP (residues 173-378 deleted) and Tcb2-ΔSMP-GFP (residues 165-369 deleted), as shown before (Toulmay and Prinz, 2012)(Fig. S1B). We then tested if localization of full-length Tcb3-GFP was altered by SMP deletions in the other tricalbins. While Tcb3-GFP was not affected by the lack of the SMP domain of either Tcb1 or Tcb2, simultaneous removal of SMP domains in both Tcb1 and Tcb2 led to a generic ER localization of Tcb3-GFP (Fig. 1C). Thus, for Tcb3-GFP to correctly localize, its own SMP domain as well as either SMP domain of Tcb1 or Tcb2 are needed. Furthermore, in cells expressing Tcb3-ΔSMP-GFP, the cortical localizations of both Tcb1-mRuby2 and Tcb2-mRuby2 were affected (Fig. S1C, D). Thus, the Tcb3 SMP domain is important for the localization of Tcb1 and Tcb2. Collectively, these results suggest that tricalbins promote each other’s localization to ER-plasma membrane contact sites through SMP domain interactions.

We hypothesized that the change in localization upon deleting the SMP domain could be underpinned by a change in the mobile behavior of tricalbin molecules. Without stable association to contact sites, tricalbins might become more dynamic and readily diffuse to other parts of the ER. To assay molecular mobility in cells, we performed fluorescence recovery after photobleaching (FRAP) experiments. We first assessed cortical Tcb3-GFP in both WT and *tcb1/2*Δ cells. We found that while the signals recovered in both cases (Fig. 1D, E), recovery was significantly faster in *tcb1/2*Δ cells (Fig. 1F, G, median half-times 12.5 s, median absolute deviation (MAD): 7.3 s, in WT cells and 3.2 s, MAD: 1.2 s, in *tcb1/2*Δ cells, P<0.0001, n≥43 FRAP events each, from 3 experimental repeats). The recovery of Tcb3-ΔSMP-GFP in WT cells and of Tcb3-GFP in Tcb1/2-ΔSMP cells was similarly fast (median half-times 4.3 s, MAD: 2.1 s, and 4.4 s, MAD: 2.1 s, respectively, P<0.0001 for both compared to WT, n≥49 FRAP events each, from 3 experimental repeats) (Fig. 1F-I). Thus, the lack of SMP domains achieves the same effect on Tcb3 dynamics as deleting the entire *TCB1* and *TCB2* genes. These results suggest that heteromerization via SMP domains is a major driver of stable accumulation of tricalbins at ER-plasma membrane contacts. This conclusion was further supported by FRAP of Tcb3-GFP in Tcb1-ΔSMP cells (Fig. S1E), which recovered with a similar half-time as Tcb3-GFP in WT cells (Fig. S1F, G). Thus, Tcb3 is more mobile only in absence of both Tcb1 and Tcb2 SMP domains, indicating redundancy in heteromerization with either Tcb1 or Tcb2. Furthermore, Tcb3-ΔSMP-GFP in *tcb1/2*1′ cells showed a recovery similar to Tcb3-GFP in *tcb1/2*Δ cells (Fig. S1F-H and 1F). Thus, the Tcb3 SMP domain alone is not sufficient to stabilize Tcb3 at the cER, indicating that putative Tcb3 homomerization through the SMP domain has no relevance for the mobile behavior of Tcb3.

### The Tcb3 C2 domains target heteromers to the cER and heteromerization is required for curvature specificity

We next explored how the C2 domains contribute to the localization of tricalbins alongside the SMP domains. We therefore investigated tricalbin mutants lacking the C-terminal part comprising all C2 domains (ΔC2) (Ikeda et al., 2020). We observed that both Tcb1-ΔC2-GFP (truncated from residue 386 onwards) and Tcb2-ΔC2-GFP (truncated from residue 378 onwards) localized to the cER exclusively (Fig. 2A), similarly to their full-length versions (Fig. S1B). In contrast, Tcb3-ΔC2-GFP (truncated from residue 491 onwards) also distributed to cytosolic ER cisternae (Fig. 2A), yet appeared mostly absent from the nuclear envelope, unlike the SMP deletion mutant (Fig. 1C). When we simultaneously imaged mRuby2-tagged Tcb1 or Tcb2 with Tcb3-ΔC2-GFP, we found that both Tcb1 and Tcb2 colocalized with Tcb3-ΔC2-GFP, hence losing their exclusive cER localization (Fig. 2B). Thus, retention at ER-plasma membrane contact sites of both Tcb1 and Tcb2 requires the C2 domains of Tcb3, but not their own C2 domains.

**Figure 2:**
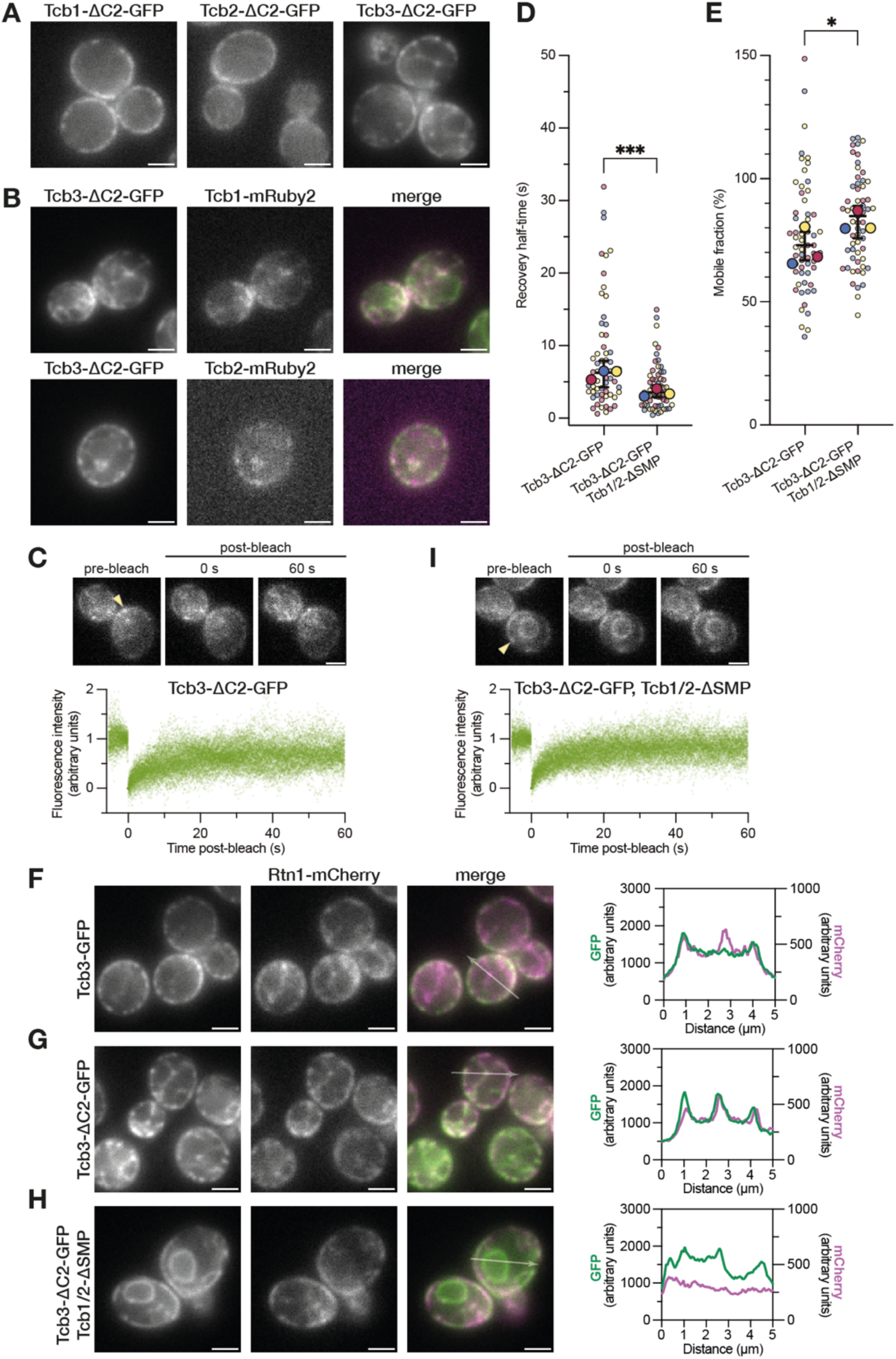
Tricalbin heteromers sense ER membrane curvature and depend on the C2 domains of only Tcb3 for targeting to the cER. **A:** Fluorescence live cell imaging of Tcb1/2/3 mutants with all C2 domains deleted. **B:** Fluorescence live imaging of cells co-expressing Tcb3-ΔC2-GFP and mRuby2 tagged Tcb1 or Tcb2. **C:** FRAP of Tcb3-ΔC2-GFP. The upper panel shows images from a representative FRAP movie, corresponding to time points before, at 0 s and 60 s after the bleaching event, indicated by an arrowhead. The lower panels show normalized and aligned fluorescence intensity measurements of all FRAP events (n=60, from 3 merged experimental repeats). **D:** Recovery half-times determined from the FRAP experiments shown in C and I. The colors encode three experimental replicates. Each data point corresponds to an analyzed FRAP event. The larger points indicate the medians of each experimental repeat, and the horizontal lines represent the median with 95% confidence interval of all data points combined. **E:** Mobile fractions determined from the FRAP experiments shown in C and I. The colors encode three experimental replicates. Each data point corresponds to an analyzed FRAP event. The larger points indicate medians of each experimental repeat, and horizontal lines represent medians with 95% confidence interval of all data points combined. **F-H**: Fluorescence live imaging of cells co-expressing Rtn1-mCherry with either Tcb3-GFP (**F**), Tcb3-ΔC2-GFP (**G**) or Tcb3-ΔC2-GFP in Tcb1/2-ΔSMP background (**H**). In the merge image of the two channels, white arrows indicate line profiles along which pixel intensities for both GFP and mCherry signals are plotted in the last panel on the right. **I:** FRAP of Tcb3-ΔC2-GFP in Tcb1/2-ΔSMP cells. The upper panel shows images from a representative FRAP movie, corresponding to time points before, at 0 s and 60 s after the bleaching event, indicated by an arrowhead. The lower panels show normalized and aligned fluorescence intensity measurements of all FRAP events (n=59, from 3 merged experimental repeats). Scale bars: 2 µm

In light of this finding and the unique localization phenotype of Tcb3-ΔC2-GFP, we performed FRAP on this mutant (Fig. 2C). Tcb3-ΔC2-GFP recovered with a median half-time of 6.3 s (MAD: 3 s) (Fig. 2D, E), which is between full-length Tcb3-GFP and the SMP deletion mutants (Fig. 1F) (n=60 FRAP events of Tcb3-ΔC2-GFP from 3 experimental repeats, P=0.0021 for comparison to Tcb3-GFP). This indicates that anchoring to the plasma membrane by the C2 domains contributes to controlling mobility of tricalbin heteromers.

The localization of Tcb3-ΔC2-GFP resembled that of reticulons, curvature inducing proteins that maintain a tubular ER network (Voeltz et al., 2006). We therefore examined its colocalization with the reticulon Rtn1-mCherry. The signal of full-length Tcb3-GFP overlaps with Rtn1-mCherry only at the cER (Toulmay and Prinz, 2012)(Fig. 2F). In contrast, Tcb3-ΔC2-GFP colocalized with Rtn1-mCherry to a high degree both cortically and internally (Fig. 2G), in line with the previous observation that tricalbins have a preference for high ER membrane curvature (Hoffmann et al., 2019). When not anchored to the plasma membrane by the C2 domains, Tcb3 thus distributes to the reticulated regions of cytosolic ER, but not to the nuclear envelope which is a low-curvature membrane devoid of reticulons. However, in Tcb1/2-ΔSMP cells, Tcb3-ΔC2-GFP became enriched in the nuclear envelope and was no longer colocalizing with Rtn1-mCherry (Fig. 2H). Thus, when heteromerization was precluded, high curvature localization was lost. By FRAP, we found that in this situation, Tcb3-ΔC2-GFP was very mobile, with a median recovery half-time of 3.6 s (MAD: 1.8 s) (Fig. 2D, E, I). This value is similar to SMP deletion mutants (Fig. 1F-I and 2D, n=59 FRAP events of Tcb3-ΔC2-GFP in Tcb1/2-ΔSMP cells, from 3 experimental repeats, P=0.06 compared to Tcb3-GFP in Tcb1/2-ΔSMP cells), indicating that mobility does not increase further when plasma membrane association is impeded in addition to heteromerization.

These results suggest that the Tcb3 C2 domains confer cortical specificity to tricalbin heteromers. Their binding to the plasma membrane restrains mobility of tricalbins to some extent. However, the restriction of lateral mobility is dominated by the heteromerization through the SMP domains, which is also required for localization to high membrane curvature.

### Intermembrane distances and mobility are coordinated through tricalbin domain architecture

Tricalbins contribute to maintaining the ER to plasma membrane proximity (Manford et al., 2012) by mediating non-specific intermembrane distances of 15-30 nm, similar to other ER-plasma membrane tethers (Collado et al., 2019; Hoffmann et al., 2019). To probe the tethering function of tricalbins, we set out to analyze a construct with modified tethering capacity. For this, we used the Tcb3_1-490_-chimeraC mutant, which we previously expressed from plasmid (Hoffmann et al., 2019) and now introduced genomically by CRISPR. In this construct, all six C2 domains are replaced with a flexible linker (GGGGS)_5_ and twice the polybasic patch by which Ist2, another ER-plasma membrane tether, binds to the plasma membrane (Fischer et al., 2009; Maass et al., 2009)(Fig. 3A). Tcb3_1-490_-chimeraC-GFP localized to cER exclusively, like WT Tcb3-GFP (Hoffmann et al., 2019). Tcb3_1-490_-chimeraC-GFP was also exclusively cortical in *tcb1/2*Δ cells and when the other major ER-plasma membrane tethers were deleted (*tcb1/2*Δ*, ist2*Δ*, scs2/22*Δ, hereafter *5*Δ for simplicity) (Fig. 3B) (Hoffmann et al., 2019; Manford et al., 2012). In contrast, WT Tcb3-GFP localization was affected in *tcb1/2*Δ cells (Fig. 1B) (Hoffmann et al., 2019), and Tcb3-GFP was barely present in *5*Δ cells (Fig. S2A, B). Thus, the chimeraC mutation renders Tcb3 localization to the cER independent of the other tricalbins.

**Figure 3:**
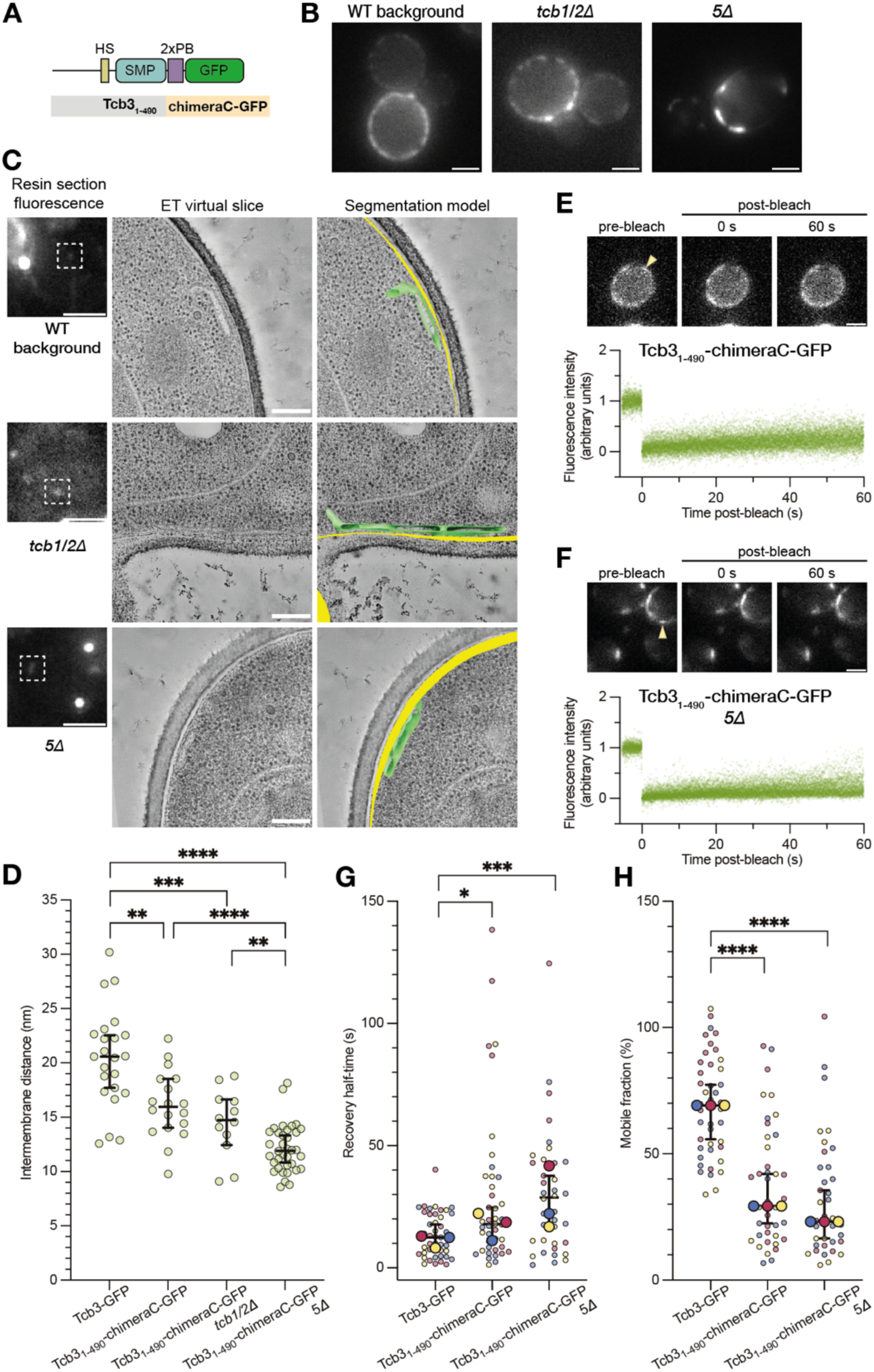
Replacing the C2 domains of Tcb3 with a linker followed by a tandem polybasic patch decreases both intermembrane distances and mobility. **A:** Schematic domain organization of the Tcb3_1-490_-chimeraC-GFP domain mutant. The 2x polybasic patch (PB) from Ist2 is preceded by 5x a GGGGS linker sequence. **B:** Live cell imaging of Tcb3_1-490_-chimeraC-GFP in WT, *tcb1/2Δ and 5Δ* genetic backgrounds. **C:** Representative CLEM images of Tcb3_1-490_-chimeraC-GFP in the same genetic backgrounds as in B. Left panels: Fluorescence microscopy of resin sections, where signals of interest are indicated by dashed squares. Middle panels: Virtual slices through corresponding electron tomograms acquired at the positions of the dashed squares. Right panels: Segmentation models, with the ER in green and plasma membrane in yellow. **D:** Intermembrane distance measurements at ER-PM contact sites mediated by WT Tcb3-GFP and Tcb3_1-490_-chimeraC-GFP in different genetic backgrounds. Horizontal lines represent medians with 95% confidence interval. **E-F:** FRAP of Tcb3_1-490_-chimeraC-GFP in the WT **(E)** and *5Δ* (F) cells. The upper panel shows images from a representative FRAP movie, corresponding to time points before, at 0 s and 60 s after the bleaching event, indicated by an arrowhead. The lower panels show normalized and aligned fluorescence intensity measurements of all FRAP events (n=41 in E, n=38 in F, from 3 merged experimental repeats). **G-H:** Recovery half-times **(G)** and mobile fractions **(H)** determined from the FRAP experiments shown in E and F, compared to WT Tcb3-GFP (same data as in Fig. 1F and G). Three experimental replicates performed on different days are color-coded, and the median for each of them denoted with a larger symbol. Horizontal lines represent medians with 95% confidence interval of all data points combined. Scale bars: 2 µm in fluorescence images, and 200 nm in electron micrographs.

We then investigated whether Tcb3_1-490_-chimeraC-GFP can change intermembrane distances at ER-plasma membrane contact sites. To this end, we employed CLEM on resin-embedded yeast cells (Kukulski et al., 2011). In electron tomograms of regions containing fluorescence signals, we analyzed the 3D membrane ultrastructure of the ER and the plasma membrane (Fig. 3C and S2C). We first acquired data on Tcb3-GFP in WT cells and measured the distance between ER and plasma membrane to be 20.6 nm (median, MAD: 2.5 nm) (Fig. 3D), similar to what we and others have reported before (Collado et al., 2019; Hoffmann et al., 2019). We then obtained measurements of Tcb3_1-490_-chimeraC-GFP in different genetic backgrounds (Fig. 3C). We found that when Tcb3_1-490_-chimeraC-GFP was expressed instead of Tcb3-GFP, the median intermembrane distance was decreased to 16.0 nm (MAD: 2.1 nm) (Fig. 3D, P=0.0011 compared to WT, n=18 Tcb3_1-490_-chimeraC-GFP and n=23 WT contact sites). In *tcb1/2*Δ cells, the median intermembrane distance for Tcb3_1-490_-chimeraC-GFP was 14.7 nm (MAD: 1.8 nm), and in *5*Δ cells it was 11.9 nm (MAD: 1.7 nm) (Fig. 3D, P=0.0002 and P<0.0001 compared to WT, respectively, n=12 in *tcb1/2*Δ cells and n=36 in *5*Δ cells). These results indicate that the Tcb3_1-490_-chimeraC-GFP mutant has an intrinsic capacity to bring the two membranes as close as 12 nm. This capacity can be hampered by the presence of other ER-plasma membrane contact site proteins, in particular its interaction partners Tcb1 and Tcb2.

We next live-imaged cells co-expressing Tcb3_1-490_-chimeraC-GFP and either Tcb1 or Tcb2 tagged with mRuby2. Both Tcb1 and Tcb2 localized exclusively to cER with their distributions matching that of Tcb3_1-490_-chimeraC-GFP (Fig. S2D). Thus, the heteromerization-dependent localization of Tcb1 and Tcb2 is maintained by the Tcb3_1-490_-chimeraC mutant. Since the localization of Tcb3_1-490_-chimeraC-GFP itself is not dependent on the other tricalbins (Fig. 3B), we analyzed its mobility in both WT and *5*Δ cells by FRAP (Fig. 3E, F). We found that 37.3% and 41.5% of FRAP events, respectively, showed no recovery, thus corresponded to areas where Tcb3_1-490_-chimeraC-GFP was completely immobile (Fig. S2E). In the remaining FRAP events, both in WT and *5*Δ cells, Tcb3_1-490_-chimeraC-GFP showed a significantly lower mobility than WT Tcb3-GFP, both in terms of recovery half-time (Fig. 3G, median in WT cells 17.9 s, MAD: 11.1 s, P=0.02 compared to Tcb3-GFP; in *5*Δ cells 28.9 s, MAD: 16.3 s, P=0.0005 compared to Tcb3-GFP, n≥38 FRAP events from 3 experimental repeats) and mobile fraction, calculated from the maximum of a fit to the fluorescence recovery curve (Fig. 3H, median in WT cells 29.3%, MAD: 13.2%, P<0.0001 compared to Tcb3-GFP, in *5*Δ cells 23.1%, MAD: 11.6%, P<0.0001 compared to Tcb3-GFP, n≥38 FRAP events each, from 3 experimental repeats). Thus, Tcb3_1-490_-chimeraC-GFP not only perturbs the contact site architecture by decreasing intermembrane distance, but it also becomes immobile. These results suggest that molecular mobility and architecture might be interdependent.

To test this idea, we attempted to further decrease the intermembrane distance using an even more truncated construct. By additionally deleting the SMP domain in Tcb3_1-490_-chimeraC-GFP, we generated Tcb3_1-272_-chimeraC-GFP (Fig. 4A), in which a sequence of only 30-40 amino acids is expected to span the cytosolic gap. In WT cells, Tcb3_1-272_-chimeraC-GFP localized exclusively to cER, but it did not efficiently recruit Tcb1-mRuby2 or Tcb2-mRuby2 to cER (Fig. S2F), in contrast to Tcb3_1-490_-chimeraC-GFP (Fig. S2D), further corroborating the necessity of SMP interactions for localization. Since the intermembrane distances mediated by Tcb3_1-490_-chimeraC-GFP were shortest in absence of other tethers, we also expressed Tcb3_1-272_-chimeraC-GFP in *5*Δ cells (Fig. 4A). Tcb3_1-272_-chimeraC-GFP localization in *5*Δ cells was not exclusively cortical, and we observed a considerable amount of protein in cytosolic ER and the nuclear envelope (Fig. 4A), unlike Tcb3_1-490_-chimeraC-GFP. To test if the impaired localization was due to the lack of the SMP domain specifically or merely due to shortening of the sequence spanning the intermembrane space, we generated a mutant in which the order of the GFP and the chimeraC mutation were reversed, hence named Tcb3_1-272_-GFP-chimeraC (Fig. 4B). In this order, the GFP could act as an inert spacer that replaces the SMP domain. We found that Tcb3_1-272_-GFP-chimeraC in *5*Δ cells had exclusively cortical localization (Fig. 4B). The cortical signals of both Tcb3_1-272_-chimeraC-GFP and Tcb3_1-272_-GFP-chimeraC corresponded to ER-plasma membrane contact sites, as confirmed by CLEM (Fig. 4C). These results suggest that the partial delocalization of Tcb3_1-272_-chimeraC-GFP was not caused by the lack of the SMP domain specifically, but rather due to the shortness of the intermembrane sequence. Indeed, we found that intermembrane distances mediated by Tcb3_1-272_-chimeraC-GFP were shorter than by Tcb3_1-272_-GFP-chimeraC (Fig. 4D, median: 10.9 nm, MAD: 0.8 nm and median: 12.5 nm, MAD 0.4 nm, respectively, P=0.0007, n=19 Tcb3_1-272_-chimeraC-GFP and n=10 Tcb3_1-272_-GFP-chimeraC contact sites). The difference of approximately 2 nm is similar to the dimensions of GFP (Yang et al., 1996). Thus, expansion of the intermembrane distance from approximately 11 nm to 13 nm, either by the SMP domain or by an inert spacer such as GFP, restores accumulation of Tcb3 constructs at cER.

**Figure 4:**
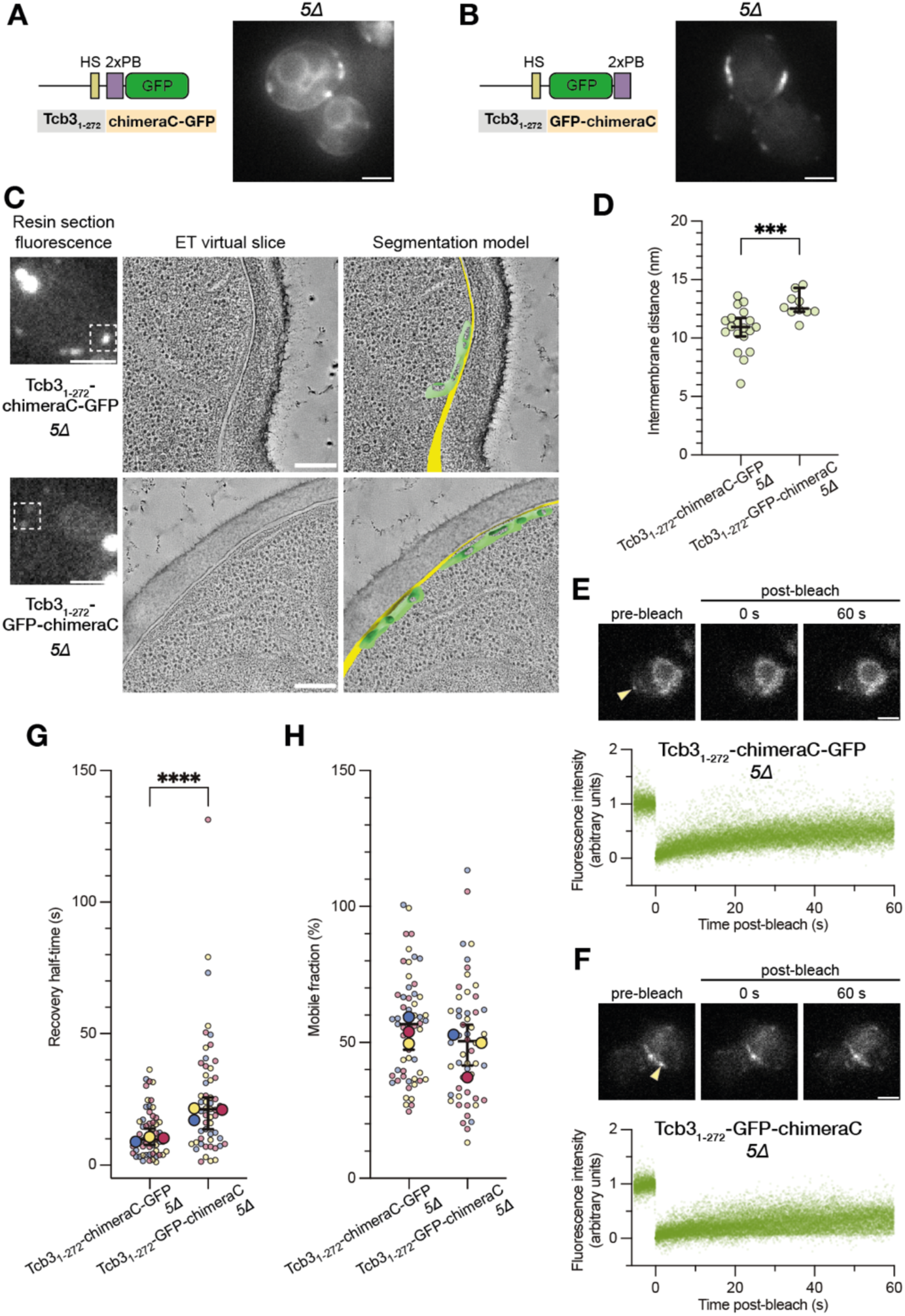
Further reducing the sequence between membranes by removing the SMP domain can impede accumulation at contact sites and increase mobility. **A:** Schematic domain organization of the Tcb3_1-272_-chimeraC-GFP mutant (left) and live cell imaging in the *5Δ* genetic background (right). **B:** Schematic domain organization of the Tcb3_1-272_-chimeraC-GFP mutant (left) and live cell imaging in the *5Δ* genetic background (right). **C:** Representative CLEM images of Tcb3_1-272_-chimeraC-GFP and Tcb3_1-272_-GFP-chimeraC, both expressed in *5Δ* cells. Left panels: Fluorescence microscopy of resin sections, where signals of interest are indicated by dashed squares. Middle panels: Virtual slices through corresponding electron tomograms acquired at the positions of the dashed squares. Right panels: Segmentation models, with the ER in green and plasma membrane in yellow. **D:** Intermembrane distance measurements at ER-PM contact sites mediated by Tcb3_1-272_-chimeraC-GFP and Tcb3_1-272_-GFP-chimeraC in *5Δ* cells. Horizontal lines represent medians with 95% confidence interval. **E-F:** FRAP of Tcb3_1-272_-chimeraC-GFP **(E)** and Tcb3_1-272_-GFP-chimeraC **(F)** in *5Δ* cells. The upper panel shows images from a representative FRAP movie, corresponding to time points before, at 0 s and 60 s after the bleaching event, indicated by an arrowhead. The lower panels show normalized and aligned fluorescence intensity measurements of all FRAP events (n=56 in E, n=54 in F, from 3 merged experimental repeats). **G-H:** Recovery half-times **(G)** and mobile fractions **(H)** determined from the FRAP experiments shown in E and F. Three experimental replicates performed on different days are color-coded, and the median for each of them denoted with a larger symbol. The horizontal lines represent the medians with 95% confidence interval of all data points combined. Scale bars: 2 µm in fluorescence images, and 200 nm in electron micrographs.

For our previous Tcb3 mutants, the two phenotypes of short intermembrane distances and generic ER distribution were associated with opposing effects on molecular mobility. Tcb3-ΔSMP-GFP with its generic ER localization was more mobile than WT Tcb3-GFP (Fig. 1), while Tcb3_1-490_-chimeraC-GFP mediated shorter distances and was less mobile than WT Tcb3-GFP (Fig. 3). Therefore, we analyzed the mobility of Tcb3_1-272_-chimeraC-GFP (Fig. 4E) and Tcb3_1-272_-GFP-chimeraC (Fig. 4F) in *5*Δ cells by FRAP. We found that Tcb3_1-272_-chimeraC-GFP recovered similarly to WT Tcb3-GFP, while Tcb3_1-272_-GFP-chimeraC showed a significantly slower mobility in comparison (Fig. 4G, H) (median half-times: 9.8 s, MAD: 5.8 s; and 21.2 s, MAD: 12.3 s, respectively, P<0.0001, n≥54 FRAP events each, from 3 experimental repeats). This is in line with our finding that the short intermembrane distances mediated by Tcb3_1-272_-chimeraC-GFP did not favor its steady accumulation and resulted in increased mobility in the ER.

### The intrinsically disordered Tcb3 N-terminus disperses Tcb3-based constructs in the ER through enhanced mobility and widens the intermembrane space

Intrinsically disordered regions (IDRs) occur frequently in contact site proteins and can act as entropic barriers to prevent crowding at membrane contact sites (Jamecna et al., 2019; Subra et al., 2023). A substantial portion of the Tcb3 sequence consists of IDRs. The cytosolically localized, N-terminal 190 amino acid residues as well as several regions between C2 domains, adding up to 274 amino acids, are likely disordered (Fig. 5A). To test if IDRs of Tcb3 play a role in contact site organization, we generated mutants in which we deleted the first 190 aa of Tcb3. Tcb3_191-1545_-GFP localized to cER in a pattern similar to WT Tcb3-GFP (Fig. S3A and 1B), indicating that the N-terminal IDR is not required for cER targeting. Next, we additionally removed the IDRs among the C2 domains through the chimeraC mutation. Tcb3_191-490_-chimeraC-GFP localized to discrete cortical puncta in both WT and *5*Δ cells (Fig. 5B). As expected for having its SMP domain, this mutant recruited both Tcb1-mRuby2 and Tcb2-mRuby2 into the puncta in WT cells (Fig. S3B, C). When the SMP domain was deleted, the resulting Tcb3_191-272_-chimeraC-GFP also localized to cortical puncta in both WT and *5*Δ cells (Fig. 5C), in line with the chimeraC mutation not depending on heteromerization for cortical localization (Fig. 3B). In contrast to Tcb3_1-272_-chimeraC-GFP (Fig. 4A), which differs only by having the N-terminal IDR, Tcb3_191-272_-chimeraC-GFP displayed no generic ER signal, meaning that all protein was efficiently targeted to contact sites when the N-terminus was removed. We wondered whether this efficient localization occurred despite a short intermembrane distance, expected due to lack of SMP domain (Fig. 4D). We therefore analyzed both Tcb3_191-490_-chimeraC-GFP and Tcb3_191-272_-chimeraC-GFP in *5*Δ cells by CLEM (Fig. 5D). We found that Tcb3_191-490_-chimeraC-GFP mediated a median intermembrane distance of 12.7 nm, MAD: 2.1 nm, while Tcb3_191-272_-chimeraC-GFP mediated a shorter median intermembrane distance of 8.4 nm, MAD: 0.7 nm (Fig. 5E, P<0.0001, n=25 Tcb3_191-490_-chimeraC-GFP and n=10 Tcb3_191-272_-chimeraC-GFP contact sites). The latter also mediated an intermembrane distance significantly shorter than when the N-terminus was present (Fig. 4C, D, P=0.0048, n=10 Tcb3_191-272_-chimeraC-GFP and n=19 Tcb3_1-272_-chimeraC-GFP contact sites). Thus, while lack of both the N-terminal IDRs and the SMP domains resulted in the shortest intermembrane distance, the protein was strictly associated with contact sites, unlike in presence of the N-terminus.

**Figure 5:**
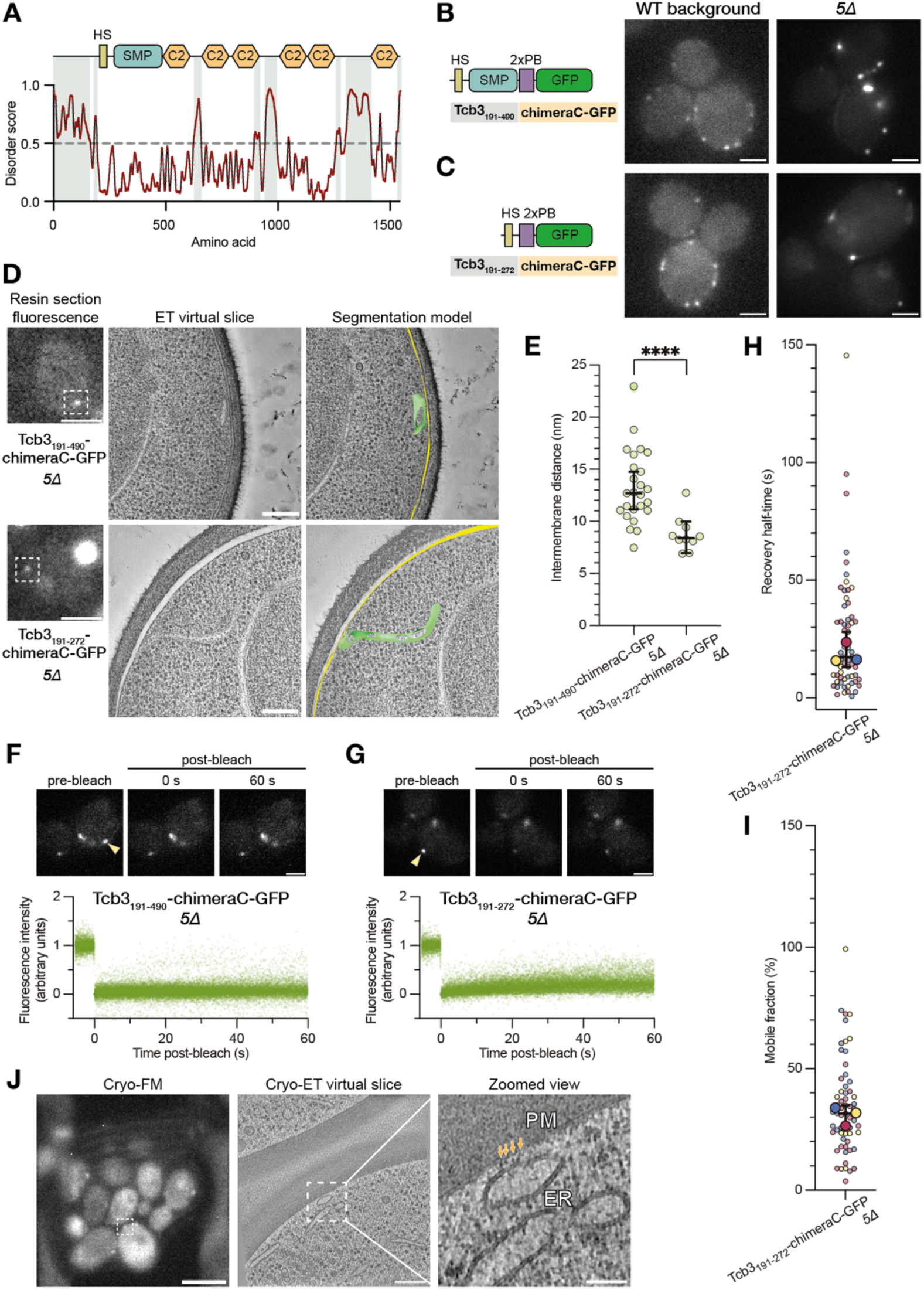
Deleting intrinsically disordered regions of Tcb3 leads to its immobile accumulation in contacts with very short intermembrane distances. **A:** Disorder prediction for the Tcb3 sequence using the Protein DisOrder prediction System PrDOS (Ishida and Kinoshita, 2007). Shaded areas mark the disordered portions of the Tcb3 sequence. **B:** Live fluorescence imaging of Tcb3_191-490_-chimeraC-GFP in WT and *5Δ* cells **C:** Live fluorescence imaging of Tcb3_191-272_-chimeraC-GFP in WT and *5Δ* cells. **D:** Representative CLEM images of Tcb3_191-490_-chimeraC-GFP (upper panel) and Tcb3_191-272_-chimeraC-GFP (lower panel) expressed in *5Δ* cells. Fluorescence microscopy of resin sections on the left, where signals of interest are indicated by dashed squares. In the middle, virtual slices through electron tomograms acquired at the fluorescent signals, and on the right segmentation models showcasing the 3D architecture of the ER-PM contact site, with the ER in green and the plasma membrane in yellow. **E:** Intermembrane distance measurements at ER-PM contact sites mediated by Tcb3_191-490_-chimeraC-GFP and Tcb3_191-272_-chimeraC-GFP in *5Δ* cells. Horizontal lines represent medians with 95% confidence interval. **F-G:** FRAP of Tcb3_191-490_-chimeraC-GFP **(F)** and Tcb3_191-272_-GFP-chimeraC **(G)** in *5Δ* cells. The upper panel shows images from a representative FRAP movie, corresponding to time points before, at 0 s and 60 s after the bleaching event, indicated by an arrowhead. The lower panels show normalized and aligned fluorescence intensity measurements of all FRAP events (n=78 in F, n=63 in G, from 3 merged experimental repeats). **H:** Recovery half-times determined from the FRAP experiments shown in G. Three experimental replicates performed on different days are color-coded, and the median for each of them denoted with a larger symbol. Horizontal lines represent medians with 95% confidence interval of all data points combined. **I:** Mobile fractions determined from the FRAP experiments shown in G. Three experimental replicates performed on different days are color-coded, and the median for each of them denoted with a larger symbol. Horizontal lines represent medians with 95% confidence interval of all data points combined. **J:** Cryo-CLEM of Tcb3_191-490_-chimeraC-GFP-mediated ER-PM contact sites in *5Δ* cells. On the left, cryo-fluorescence microscopy of a cryo-FIB milled lamella. The signal of interest is indicated by the white dashed square. In the middle, virtual slice through a cryo-electron tomogram acquired at the fluorescent signal. On the right, zoomed view of a Tcb3_191-490_-chimeraC-GFP-mediated ER-PM contact site. Bridging densities between membranes are indicated with orange arrows. Scale bars: 2 µm in B, C, D (left panels; fluorescence), F and G; 200 nm in D (middle panel, ET images) and J (middle panel, ET image); 5 µm in J (left panel, cryo-fluorescence); 50 nm in J (right panel, zoom-in).

When we performed FRAP on these two mutants in *5Δ* cells, we found no detectable recovery for Tcb3_191-490_-chimeraC-GFP, and practically all the molecules in the puncta were immobile (Fig. 5F). Tcb3_191-272_-chimeraC-GFP recovered, albeit slowly with a median half-time of 17.2 s (MAD: 12.1 s) and a median mobile fraction of 31.6 % (MAD: 9 %) (Fig. 5G-I, n=63 FRAP events from 3 experimental repeats). Thus, removal of the N-terminal IDRs clusters the protein in contact sites and disrupts mobility entirely. Additional removal of the SMP domain renders the intermembrane distances exceptionally short and restores minimal mobility. Despite the regained mobility, without the IDRs the protein does not distribute to non-cortical parts of the ER.

The 4.3 nm difference in intermembrane distance between Tcb3_191-490_-chimeraC-GFP and Tcb3_191-272_-chimeraC-GFP matched the length of an SMP domain (Schauder et al., 2014). This suggested that single SMP domains of Tcb3_191-490_-chimeraC-GFP could be arranged perpendicularly to the membranes, an orientation previously suggested for full-length tricalbins (Hoffmann et al., 2019; Wang et al., 2023). Such a structural integrity is not necessarily expected for this construct, given its lack of mobility, which could be interpreted as a structural collapse. We therefore decided to investigate the arrangement of Tcb3_191-490_-chimeraC-GFP relative to the membranes in *5*Δ cells by cryo-ET, targeted by cryo-correlative microscopy (Wozny et al., 2023). At ER-plasma membrane contact sites marked by the signal of Tcb3_191-490_-chimeraC-GFP, in regions of 10-13 nm intermembrane distance, we mostly observed densely packed bridging structures (Fig. 5J and S4A, B). In some cases, we observed a continuous layer of density (57% and 43%, respectively, N=44 contact sites) (Fig. S4B, C). These results suggest that a significant fraction of the SMP domains was likely in an orientation parallel to the membranes. However, most SMP domains of Tcb3_191-490_-chimeraC-GFP maintained an arrangement perpendicular to the membranes. This observation indicates that the structural integrity of tricalbins displays a considerable degree of robustness to perturbations such as impaired mobility and intermembrane distances.

## Discussion

### Tcb3 is the master organizer of tricalbin heteromers at the cER

In this study, we dissected how the domain organization of tricalbins drives and fine-tunes their function as ER-plasma membrane tethering proteins. We found that a key property of tricalbins is their association into heteromers via their SMP domains. Tcb3 interacts with both Tcb1 and Tcb2 via the SMP domains, and this heteromeric interaction is essential for the localization of tricalbins to cER. Heteromerization is also required for localization to high membrane curvature ER. Previous work showed that SMP domains target tricalbins to cER (Toulmay and Prinz, 2012), and that the transmembrane domain is responsible for curvature localization (Hoffmann et al., 2019). Our present results show that the basis of both these functions is heteromerization, without which neither can occur. A consequence of heteromerization is a substantial slowdown of mobility within the ER membrane. This slowdown contributes to retention of tricalbin heteromers at ER-PM contact sites, which requires the C2 domains of Tcb3, while the C2 domains of Tcb1 and Tcb2 are not sufficient. Previously, yeast two-hybrid experiments have reported that Tcb2 interacts with both Tcb1 and Tcb3 through the C2 domains (Creutz et al., 2004). Our results do not exclude an interaction between C2 domains, which could additionally strengthen the anchoring to the plasma membrane or serve a regulatory function. Another heteromerization interface in the hydrophobic stretch containing the transmembrane domains (Hanaoka et al., 2024) is also compatible with our results, and could be supporting curvature localization. In fact, our AlphaFold3 based heterodimeric models of Tricalbin constructs predict interfaces between both the SMP and the hairpin structure of the hydrophobic stretch. Nevertheless, our experiments indicate that heteromerization is dominated by the SMP domain interaction, and none of the other interfaces are sufficient for the tricalbins to recruit each other to ER-plasma membrane contact sites. SMP domains have been suggested to also homomerize, but curiously, our results suggest that homomerization of Tcb3 SMP domains, if it occurs, is not relevant for cER and high-curvature localization. This could point towards an additional role for heteromerization via the C2 domains or hairpin domains of Tcb1 and Tcb2, for instance to assemble higher-order oligomers, similarly to reticulons (Shibata et al., 2008). The SMP domains likely form head-to-head dimers, as in structures of their mammalian orthologues the extended synaptotagmins (Schauder et al., 2014) and predicted by AlphaFold3 (Abramson et al., 2024), which might be the prerequisite for further interactions.

Regardless of the exact oligomeric state, our results suggest that the tricalbins rely on each other’s structural elements to form assemblies that influence their localization and retention. We found Tcb1 and Tcb2 to be redundant regarding their tethering functions. If heteromerization between them occurs, it is unable to replace heteromerization with Tcb3, as in absence of the Tcb3 SMP domain both Tcb1 and 2 lose specificity for cER, despite the other partner being available. Similarly, they are unable to rescue the lack of the Tcb3 C2 domains through their respective C2 domains. Furthermore, while the master organizer Tcb3 requires presence of one of the other two, it does not seem to matter which one. This finding is consistent with a high sequence similarity between Tcb1 and Tcb2 (55% identity, in contrast to 26% for either Tcb1 or Tcb2 compared to Tcb3, according to Clustal2.1 (Madeira et al., 2024)).

### Intermembrane distance and lateral mobility of tricalbins are fine-tuned

Our study also addresses the relationship between intermembrane distance and protein accommodation. We generated chimeric Tcb3 variants, in which we replaced the C2 domains by the polybasic patch of Ist2. These constructs imposed shorter intermembrane distances and displayed impaired mobility. Previous work had shown that by itself, the tandem polybasic patch of Ist2 is highly mobile at the plasma membrane, as in FRAP experiments its signal recovers fully within just a few seconds (Sugiyama and Kono, 2024). Therefore, the changes in the mobility of chimeraC mutants are likely not due to the polybasic patch, but rather due to the effect the mutation has on the intermembrane space. Shortened intermembrane distances slow down the lateral diffusion of tricalbins. However, at very short intermembrane distances, mediated by linker sequences of 30-40 amino acids, movement is to some extent restored, and tricalbin constructs “leak out” of ER-plasma membrane contact sites. Thus, such narrow spacing between the membranes appears to be unfavorable for stable accommodation, for example for electrostatics or steric reasons. The latter is supported by two of our experimental results. First, when we increase the intermembrane linker sequence by placing a GFP moiety, intermembrane distances are larger, and leakage is prevented. Second, by comparing constructs with and without the N-terminal IDRs, we found that presence of the IDR widens the intermembrane distance for the construct with the shortest intermembrane linker. In addition, the IDR laterally disperses the Tcb3 construct within the cER by increasing its mobility. This is likely linked to an enlarged hydrodynamic radius, as previously shown for OSBP (Jamecna et al., 2019). In absence of the N-terminal IDR, the Tcb3 construct remains confined to ER-plasma membrane contact sites, despite the narrow intermembrane space.

Our results show that the intermembrane distances and the mobility of Tcb3 can be modulated by the protein domain architecture. In ER-PM contact sites mediated by WT Tcb3, the intermembrane spaces are considerably wider than we show would be possible. The protein mobility is moderate, yet constrained by the heteromerization. Such a membrane contact site environment might be ideal to accommodate various molecular functions, including lipid transfer, and at the same time allow sufficient lateral movement of the populating proteins. For instance, mobility and turnover of tricalbins might be important for their function in sustaining rapid lipid fluxes for plasma membrane repair or expansion. Although tricalbins are not essential for cell fitness under normal growth conditions, their lipid transfer function becomes important in response to heat shock (Collado et al., 2019; Thomas et al., 2022) and hypoosmotic swelling (Mu et al., 2024; Smith et al., 2024). The intermembrane distances mediated by WT tricalbins are similar to those mediated by the other ER-plasma membrane tethers, as well as those reported for other organelle contacts (Collado et al., 2019; Fernandez-Busnadiego et al., 2015; Hoffmann et al., 2019), and could thus represent a compromise between proximity and separation. For example, the ER-resident phosphatase Sac1 which hydrolyses PI4P both in the ER (trans) and the plasma membrane (cis) may be controlled by intermembrane distance (Manford et al., 2010; Stefan et al., 2011). Decreasing the distance increases accessibility of plasma membrane PI4P to Sac1 for its cis-activity, leading to dissipation of the functionally important PI4P gradient between ER and plasma membrane (Zewe et al., 2018). Conversely, an increased intermembrane distance might hamper lipid transfer between the membranes, as shown for SMP domains *in vitro* (Bian et al., 2019).

Our results also indicate that tricalbin architecture can be sustained in varying contact site environments. This is particularly supported by our cryo-ET data, which shows that the SMP domain arrangement can be maintained despite the contact site architecture, protein content and dynamics being severely affected. Such robustness could be important for continued tethering in fluctuating conditions, especially in situations of stress.

In summary, we suggest that the tethering function of tricalbins intricately combines structural control and fine-tuned flexibility for regulating intermembrane spacing, membrane ultrastructure and lateral protein mobility at ER-plasma membrane contact sites.

## Materials and methods

### Yeast strains construction

A list of yeast strains presented in this study is provided as Supplementary Table 1. Endogenous tagging and gene deletions in *Saccharomyces cerevisiae* S288C were performed using cassettes amplified from plasmids (Janke et al., 2004): pFA6a-eGFP-HIS3MX6, pFA6a-mCherry-kanMX4. pFA6a-yomRuby2-KanMX4 (Hoffmann et al., 2019), pFA6a-hphNT1 and pFA6a-klURA3. Genomic mutations Tcb1-1′SMP, Tcb2-1′SMP, Tcb3-1′SMP, Tcb1-1′C2, Tcb2-1′C2, Tcb3-1′C2, Tcb3_1-490_-chimeraC, Tcb3_1-272_-chimeraC, Tcb3_1-272_-GFP-chimeraC and Tcb3_191-1545_ were generated using CRISPR editing according to (Laughery et al., 2015). Oligonucleotide pairs encoding the 20 nt sgRNA sequence in the format GATCxxxxxxxxxxxxxxxxxxxxGTTTTAGAGCTAG and CTAGCTCTAAAACxxxxxxxxxxxxxxxxxxxx for the reverse complement (Supplementary Table 2) were hybridized and ligated into SwaI and BclI linearised plasmid pML107 and cloned in TOP10 cells. Templates for homology directed repair (HDR) were PCR-amplified to have at least 30 nucleotide long homology arms (Supplementary Table 3).

### Live cell imaging

Liquid cultures of yeast cells were grown in synthetic complete medium without tryptophan, supplemented with 2% glucose (SC-Trp) at 25 °C for all live imaging experiments. Cells in mid-log growth phase were mounted on concanavalin A-coated 1.5 glass coverslips and imaged on a widefield Nikon Eclipse Ti2 microscope using a CFI Apochromat TIRF 100×/1.49 NA oil objective and controlled by the NIS-Elements software. A photometrics Prime BSI sCMOS camera was used for collecting equatorial plane images. The samples were illuminated with 470 nm and 555 nm excitation lines provided by the Lumencor SpectraX LED light source (Chroma), with quad band filter set 89000 ET Sedat Quad (Chroma) including ET490/20x, 89100bs, ET525/36m for green fluorescence and ET555/25x, 89100bs, ET605/52m for red fluorescence respectively. An emission filter wheel (Nikon) was set to 535 nm for green fluorescence and 638 nm for red fluorescence. LED light intensities were set to 10% and exposure times ranged between 0.5 s and 5 s for acquiring images. The software Fiji (Schindelin et al., 2012) was used for preparing figures.

### Fluorescence recovery after photobleaching (FRAP)

FRAP experiments were carried out on an Olympus IX83 widefield microscope equipped with a 150×/NA1.45 objective and ImageEM X2 EM-CCD camera (Hamamatsu), controlled by the VisiView software (Visitron Systems). Widefield equatorial plane movies of yeast cells expressing GFP-tagged proteins were recorded using the TIRF 488 nm laser line at 0°, controlled by iLas2 software (Roper Scientific). Excitation and emission were filtered through a TRF89902 405/488/561/647 nm quad-band filter set (Chroma). Exposure time was set to 100 ms with no intervals, and the EMCCD gain at 255. Photobleaching was achieved with the FRAP488 laser controlled by the iLas2 software with the intensity 100-120 ms/pixel.

The FRAP data was analyzed using the Frapit toolbox (https://github.com/apicco/frap). The movies were manually cropped in Fiji and slices with photo-bleach events were deleted. The cropped movies were then processed using the BGN_FRAP_V1.ijm script (https://github.com/apicco/frap/blob/master/BGN_FRAP_V1.ijm available on GitHub). The script subtracts background using the rolling ball algorithm (radius = 90 pixels) and corrects for photobleaching based on median filtering (radius = 8 pixels). Bleached regions of interest were marked with a 5 pixel thick line profile and kymographs generated with the MultiKymograph function in Fiji.

Pixel intensities were extracted from kymographs and recovery parameters measured with a script in the Google Collaboratory (https://bit.ly/analyse-your-frap). The data was full-range normalized to 10 frames before the bleach event. Intervals between frames were calculated based on time stamps in the movie. Their average was determined to be 137 ms, accounting for a 37 ms delay in the acquisition, and was used in the analysis. Normalized recovery was fitted according to the exponential equation: *F_FRAP_*(*t*) = *A*(1 − *e*−*^kt^*), where k = ln2/τ

A represents the mobile fraction and τ the recovery half-time. Both mobile fractions and the respective half-times were analyzed for all strains. Occasionally, some FRAP events were fitted with mobile fractions significantly higher than 100%, likely due to axial or lateral movements during acquisition and were thus excluded from the analysis. Noisy bleach events or those occurring at the end of movies which could not be reliably fitted were also excluded.

### Correlative light and electron microscopy (CLEM)

Room-temperature CLEM was based on (Kukulski et al., 2012) with some adaptations. Yeast strains expressing a GFP-tagged protein of interest were grown in SC-Trp media at 25 °C, as described in the live imaging section. In mid-log phase, the cells were collected by vacuum filtration and vitrified with a Leica EMPACT-2 high pressure freezer in in 200 µm deep wells of 3 mm gold-coated copper membrane carriers (Wohlwend GmbH), coated with 1% lecithin. The resulting cryo-immobilized samples were then processed by freeze substitution, first shaking on dry-ice for 2 to 3 h in glass-distilled acetone with 0.05% (w/v) uranyl acetate (McDonald and Webb, 2011), before being placed in a Leica EM AFS2, where freeze substitution continued for another 16 h at −90 °C. The temperature was subsequently raised to −45 °C (5 °C/h) and the samples washed with acetone three times, 10 min each. This was followed by gradual infiltration with Lowicryl HM20 (Polysciences) resin diluted in acetone at the following concentrations (10%, 25%, 50%, 75%, 4 h each), while raising the temperature to −25 °C. After that, three exchanges of 100% Lowicryl 10 h each were carried out and the resin polymerized by UV at −25 °C for 48 h. The temperature was finally brought up to 20 °C (5 °C/h) and UV polymerization continued for 48 h.

Embedded cells were sectioned on a Ultracut E ultramicrotome (Reichert-Jung) equipped with a DiATOME ultra 45° diamond knife. The resulting 200 nm thin sections were applied to carbon-coated 200 mesh copper EM grids (Agar Scientific) and 0.1 μm TetraSpeck fluorescent microspheres (Invitrogen), diluted in PBS, immediately adsorbed. Fluorescence imaging of resin sections was performed on the same microscopic set-up as described in the live imaging section. Due to uneven section deposition on the EM grid, z-stacks with 0.1 or 0.2 μm step size were acquired to maximize capturing as many target regions in focus as possible.

For electron tomography, 15 nm protein A-coupled gold beads (Aurion) were applied to the grids as fiducial markers to aid reconstruction, followed by post-staining with Reynolds lead citrate (Reynolds, 1963) for contrast enhancement. Tilt-series were acquired using a Model 2020 (Fischione Instruments) holder inserted into a FEI Tecnai Spirit TEM operated at 80 kV with a Tungsten filament. Digital images of the sample were recorded using SerialEM (Mastronarde, 2005) on a FEI 4k Eagle camera in dual-axis mode (Mastronarde, 1997) over a tilt range from −55° to 55° with 1° increment and a pixel size of 1.2 nm. Additionally, lower magnification montages at 3.3 nm pixel size were collected for correlation to fluorescence images. The IMOD software package was used to reconstruct tomograms (Kremer et al., 1996). TetraSpeck-based correlation was performed using MATLAB-based scripts described in (Kukulski et al., 2011).

### Intermembrane distance analysis

ER-plasma membrane contact sites correlated to a fluorescence signal were subjected to intermembrane distance analysis. Both the ER membrane facing the plasma membrane and the plasma membrane were parametrized by manually defining points through the tomographic volume approximately every 6 nm along the z axis. Using MATLAB scripts described in (Ganeva et al., 2023), surfaces were rendered through these points, modelling the apposing ER and plasma membrane bilayers. The distances between them were measured over a grid defined by 5 nm spacing. For each contact site, the extracted distances were analyzed in R using the ggplot2 package (Wickham, 2016). A probability density function was fitted through the data specifying a bandwidth of 5 nm. The peak value of the resulting plot was taken as the final measurement of intermembrane distance for each contact site. In total, 10-40 intermembrane distance measurements were made for each yeast strain. In a few cases with large, complex ER morphologies, the contact area was split into two or three separately measured areas. Three dimensional segmentations of representative ER-plasma membrane contact sites were manually drawn in IMOD using the sculpt tool and rendered in UCSF ChimeraX (Meng et al., 2023).

### Vitrification for cryo-ET

Cells expressing Tcb3_191-490_-chimeraC-GFP in the *5Δ* genetic background were grown to exponential mid-log phase in the SC-Trp medium at 25 °C. 6 ml of culture were pelleted and resuspended in 2 ml SC-Trp supplemented with 15% dextran (Sigma, 40 kDa Mw) as a cryo-protectant. Of this culture, 5 μl were applied to Quantifoil R2/2 Cu 200 mesh grids, glow discharged at 15 mA for 30 s before excess solution was blotted away manually with a Whatman filter paper (No. 1) on the back side of the grid for 14 seconds, before plunge-freezing using a custom-made manual plunger with a temperature-controlled liquid ethane cup set to -181 °C (Russo et al., 2016).

### Cryo-focused ion beam (cryo-FIB) milling and integrated fluorescence light microscopy (iFLM)

Plunge-frozen grids were clipped into AutoGrids (ThermoFisher Scientific) and mounted onto the 35° shuttle for the Aquilos 2 Cryo-FIB (ThermoFisher Scientific), equipped with an integrated fluorescence light microscope (iFLM). Grid overviews were acquired as SEM tile scans in the MAPS software (ThermoFisher Scientific), from which milling targets could be identified. The samples were sputter coated with a thin conductive layer of platinum. Subsequently, gas injection system (GIS) coating was applied for 1 min 30 s or 2 min to deposit a thick, protective organometallic platinum layer. FIB-milling was automated with the AutoTEM Cryo software (ThermoFisher Scientific) where relevant parameters including ion beam currents as well as milling patterns and lamella dimensions were defined. Eucentric heights and milling angles for targets chosen in the MAPS software were automatically calculated by AutoTEM Cryo. Automated FIB-milling was performed using standard procedures (Wagner et al., 2020). First, micro expansion joints (Wolff et al., 2019) were cut out on both sides of the lamella. In subsequent steps, the milling currents were decreased (1.0 nA, 0.3 nA and finally 0.1 nA) to obtain approximately 1 μm thick lamella. Fluorescence z-stacks of the resulting lamellae were acquired using the iFLM software in the green channel with 1 s exposure time and excitation light intensity set to 30%. Only lamellae containing signals of interest were next manually polished with ion-beam currents of 50 and 30 pA. The fine-milled lamellae were imaged once more by iFLM.

### Cryo-electron tomography (cryo-ET)

Cryo-FIB-milled lamellae were imaged on a Krios G4 C-FEG cryo-TEM operated at 300 kV equipped with a Falcon 4i detector used in counting mode and a Selectris X energy filter. Tilt series were acquired using Tomography 5 Software (ThermoFisher Scientific). Medium magnification maps of the lamellae were acquired at a pixel size of 3.8 nm. These served to assess the quality of the lamellae and identify ER-plasma membrane contact sites of interest by visual correlation to the iFLM images. Tilt series were recorded using a grouped-dose symmetric tilt scheme (Hagen et al., 2017), starting at the pre-tilt angle with 1° increments in alternating groups of four tilt steps up to ±56°. The energy filter slit width was set to 20 eV. The dose rate was 7.5 - 9 e^−^/px/s and the targeted dose per tilt image ranged between 1.0 - 1.2 e^−^/Å^2^ fractioned over 6 or 8 frames. Nominal defocus was set to either -3 µm or -5 µm. The calibrated pixel size was 2.97 Å. Tomograms were reconstructed with IMOD (Kremer et al., 1996) at bin 2, aligning the tilt series with patch tacking and using the Simultaneous Iterative Reconstruction Technique (SIRT). For figure panels, a median filter was applied to improve visibility.

### Statistical analysis

Statistical tests to compare intermembrane distances, FRAP recovery half-times and mobile fractions were performed using GraphPad Prism. All FRAP experiments were done in three independent experimental repeats, each performed on a separate day using a fresh liquid culture of the corresponding yeast strains. The three experimental repeats are depicted by the superplots in the figure panels (Lord et al., 2020). Statistical tests were performed using pooled data points from all three experimental repeats. For all comparisons, non-parametric Mann-Whitney tests were used to account for non-normal distributions of data points. Statistical significance is indicated in figure panels as follows: P σ 0.05 as *, P σ 0.01 as **, P σ 0.001 as ***, p σ 0.0001 as ****. Exact P values are indicated in the text.

## Acknowledgements

We thank the Microscopy Imaging Center (MIC) of the University of Bern and the Dubochet Center for Imaging (DCI) Bern for access to microscopes and support in data acquisition. We thank Patrick Hoffmann for generating yeast strains used in this study. We are grateful to Daniel Lévy and Manuela Dezi for advice and discussions, and to Kukulski group members for discussions and comments on the manuscript. This work was supported by Swiss National Science Foundation (SNSF) project grant 201158 to W.K., NCCR TransCure grant 185544 and SNSF project grant 212288 to M.K..

## Declaration of interests

The authors declare no competing interests.

---

**Supplementary Figure S1:**
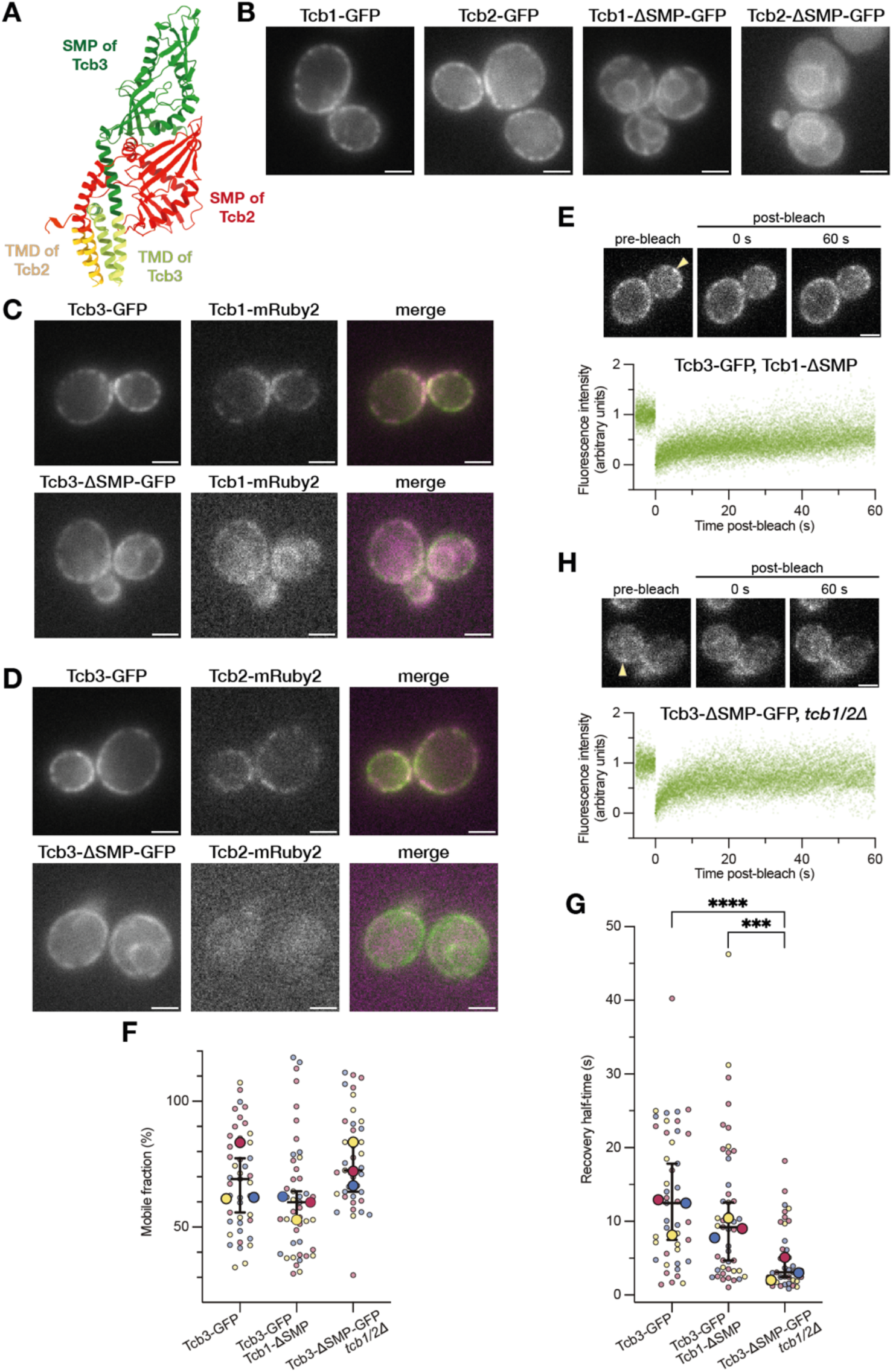
Cortical colocalization between tricalbins depends on the SMP domains. **A:** AlphaFold3 (Abramson et al., 2024) prediction of a complex between Tcb3 (green) and Tcb2 (red), using the sequences containing the helical hairpins with the hydrophobic stretch plus the SMP domains (residues 191-490 of Tcb3, and residues 91-371 of Tcb2). Residues predicted as transmembrane regions using TMHMM 2.0 (Krogh et al., 2001) are shown in light green (for Tcb3) and in light orange (for Tcb2). **B:** Fluorescence live imaging of cells expressing GFP-tagged Tcb1 and Tcb2 and the respective SMP domain mutants. **C:** Fluorescence live imaging of cells co-expressing Tcb3-GFP (upper panel) or Tcb3-ΔSMP-GFP (lower panel) together with Tcb1-mRuby2. **D:** Fluorescence live imaging of cells co-expressing Tcb3-GFP (upper panel) or Tcb3-ΔSMP-GFP (lower panel) together with Tcb2-mRuby2. **E:** FRAP of Tcb3-GFP in Tcb1-ΔSMP cells. The upper panel shows images from a representative FRAP movie, corresponding to time points before, at 0 s and 60 s after the bleaching event, indicated by an arrowhead. The lower panels show normalized and aligned fluorescence intensity measurements of all FRAP events (n=44, from 3 merged experimental repeats). Three experimental replicates performed on different days are color-coded, and the median for each of them denoted with a larger symbol. The horizontal lines represent the medians with 95% confidence interval of all data points combined. **F:** Mobile fractions determined from the FRAP experiments shown in D and G together with the ones from Fig. 1D. **G:** Recovery half-times determined from the FRAP experiments shown in D and G together with the ones in Fig.1D. Three experimental replicates performed on different days are color-coded, and the median for each of them denoted with a larger symbol. The horizontal lines represent the medians with 95% confidence interval of all data points combined. **H:** FRAP of Tcb3-ΔSMP-GFP in *tcb1/2Δ* cells. The upper panel shows images from a representative FRAP movie, corresponding to time points before, at 0 s and 60 s after the bleaching event, indicated by an arrowhead. The lower panels show normalized and aligned fluorescence intensity measurements of all FRAP events (n=38, from 3 experimental repeats). Scale bars: 2 µm

**Supplementary Figure S2:**
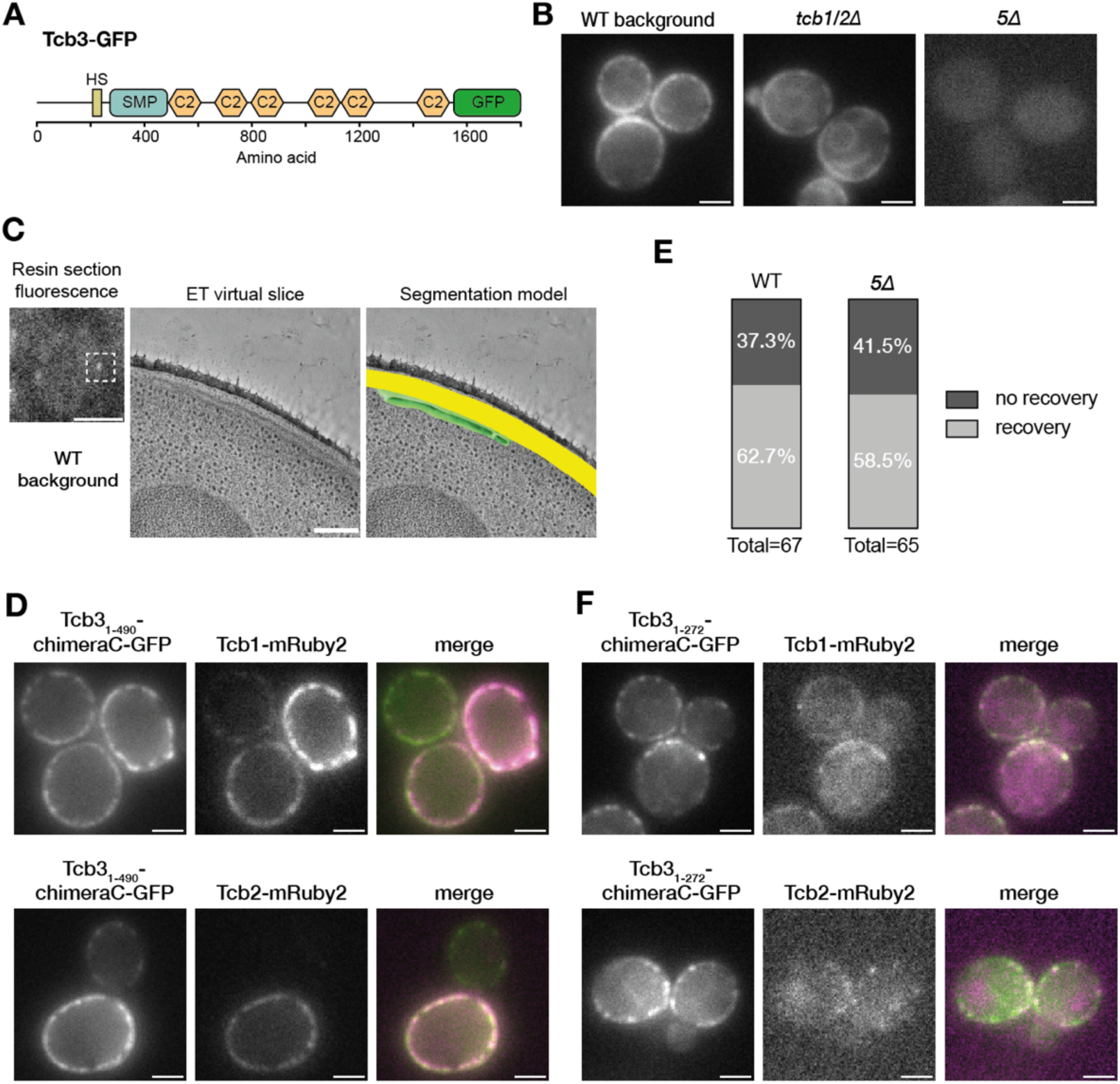
The chimeraC mutation does not prevent colocalization of tricalbins unless the SMP domain is also lacking. **A:** Domain organization schematic for Tcb3-GFP **B:** Fluorescence live imaging of cells expressing Tcb3-GFP in different backgrounds: WT*, tcb1/2Δ* (a different example of the same strain is shown in Fig. 1B), and *5Δ* cells. **C:** Representative CLEM images of Tcb3-GFP expressing cells. Fluorescence microscopy of a resin section on the left, where a signal of interest is indicated by a dashed square. In the middle, a virtual slice through an electron tomogram acquired at the fluorescent signal, and on the right a segmentation model showcasing the three-dimensional architecture of the ER-PM contact site. **D:** Fluorescence live imaging of cells co-expressing Tcb3_1-490_-chimeraC-GFP together with Tcb1-mRuby2 (upper panel) or Tcb2-mRuby2 (lower panel). **E:** Bar graphs showing percentages of Tcb3_1-490_-chimeraC-GFP FRAP events with or without measurable recovery. **F:** Fluorescence live imaging of cells co-expressing Tcb3_1-272_-chimeraC-GFP together with Tcb1-mRuby2 (upper panel) or Tcb2-mRuby2 (lower panel). Scale bars: 2 µm in all fluorescence images, and 200 nm in electron micrographs.

**Supplementary Figure S3:**
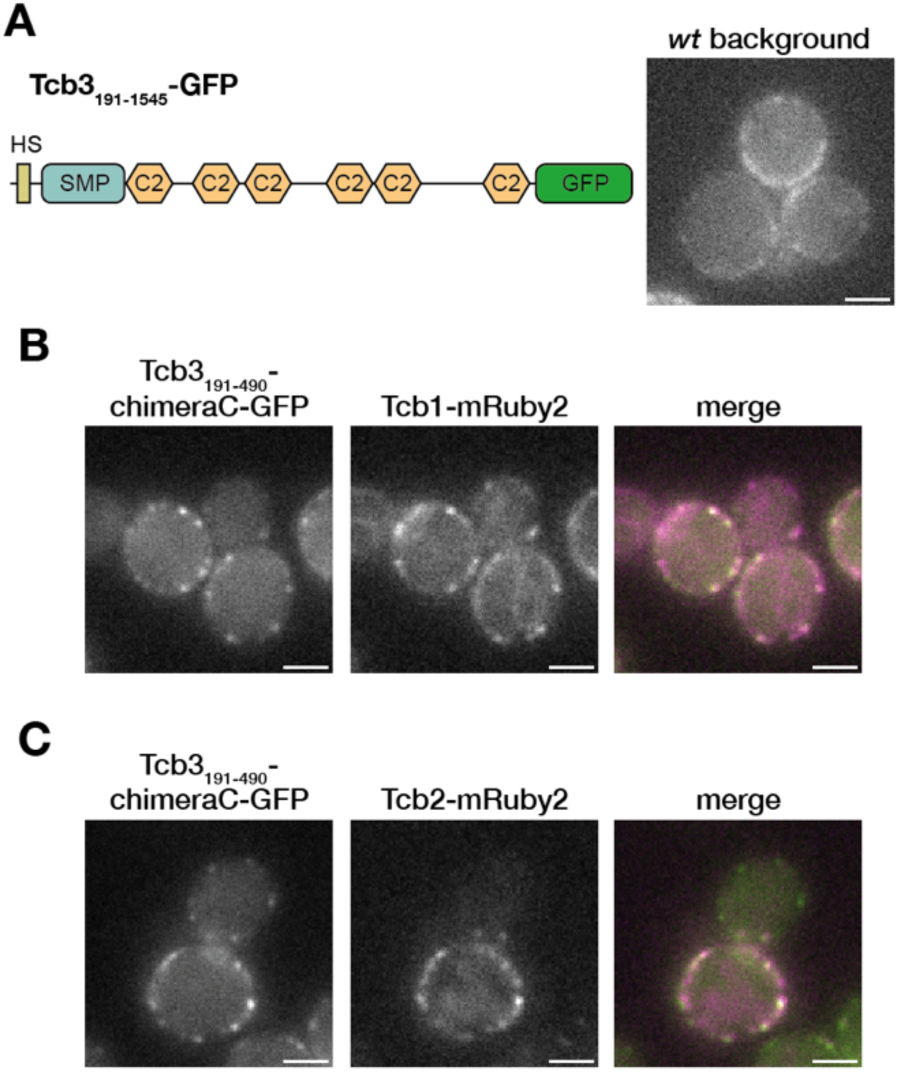
Tcb1 and Tcb2 can be recruited into the Tcb3_1-490_-chimeraC puncta. **A:** Domain organization of Tcb3_191-1545_-GFP and fluorescence live imaging of Tcb3_191-1545_-GFP in WT cells. **B:** Fluorescence live imaging of cells co-expressing Tcb3_191-490_-chimeraC-GFP together with Tcb1-mRuby2. **C:** Fluorescence live imaging of cells co-expressing Tcb3_191-490_-chimeraC-GFP together with Tcb2-mRuby2. Scale bars: 2 µm.

**Supplementary Figure S4:**
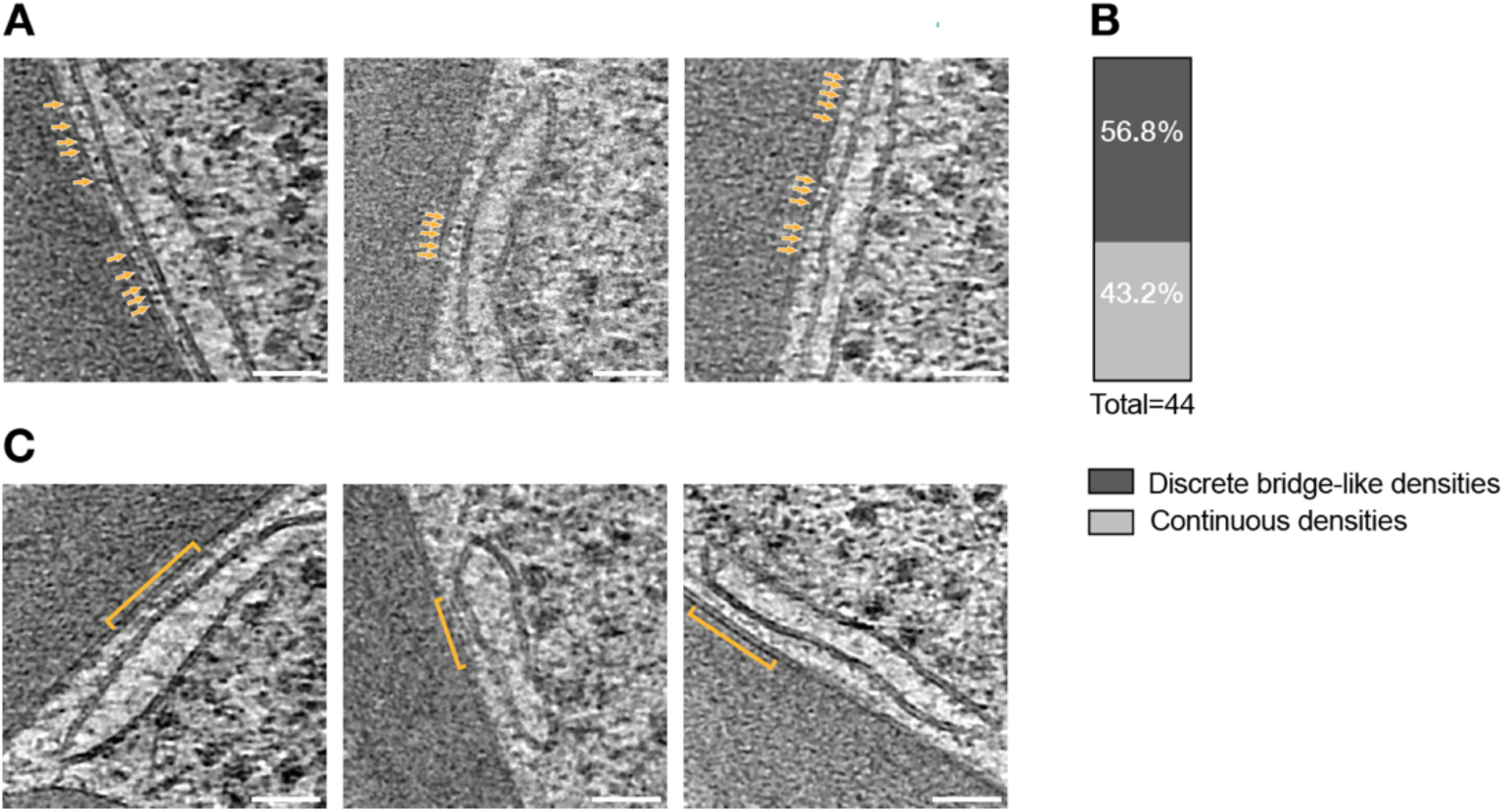
Tcb3_191-490_-chimeraC-GFP-mediated ER-plasma membrane contact sites consist of structures bridging the intermembrane space. **A:** Cryo-ET of Tcb3_191-490_-chimeraC-GFP mediated ER-plasma membrane contact sites in *5Δ* cells revealing distinct bridge-like structures perpendicular to the bilayers, indicated by orange arrowheads. **B:** Percentage analysis for the two contact site classes shown in A and C. **C:** Cryo-ET of Tcb3_191-490_-chimeraC-GFP mediated ER-plasma membrane contact sites in *5Δ* cells showing a continuous density layer lining the cytosolic gap, indicated by the orange arrowheads. Scale bars: 50 nm.

## Notes

### Competing Interest Statement

The authors have declared no competing interest.

