## Supplementary Tables for "The molecular basis of tricalbin-mediated membrane contact site organization in cells"

[1]Institute of Biochemistry and Molecular Medicine, University of Bern, 3012 Bern, Switzerland
[2]Graduate School for Cellular and Biomedical Sciences, University of Bern, 3012 Bern, Switzerland
[3]Department of Biochemistry, University of Geneva, 1211 Geneva, Switzerland
[4]current affiliation: MIDA.science, 1200 Geneva, Switzerland

**Supplementary Table 1: Yeast strains**

| Strain | Genotype | Figures | Reference |
|---|---|---|---|
| WKY0117 | *MAT$\alpha$, his3-$\Delta$200, leu2-3,112, ura3-52, lys2-801, TCB3-EGFP::HIS3MX6* | (1B,D,F,G), (2D,G,H), (S2B-C) | Hoffmann *et al.* 2019 |
| WKY0147 | *MAT$\alpha$, his3-$\Delta$200, leu2-3,112, ura3-52, lys2-801, TCB1-EGFP::HIS3MX6* | (S1B) | Lab stock |
| WKY0149 | *MAT$\alpha$, his3-$\Delta$200, leu2-3,112, ura3-52, lys2-801, TCB2-EGFP::HIS3MX6* | (S1B) | Lab stock |
| WKY0204 | *MAT$\alpha$, his3-$\Delta$200, leu2-3,112, ura3-52, lys2-801, TCB1$\Delta$::URA3, TCB2$\Delta$::hphNT1, TCB3-EGFP::HIS3MX6* | (1B, E-G), (S2B) | Lab stock |
| WKY0270 | *MATa, his3-$\Delta$200, leu2-3,112, ura3-52, lys2-801, SCS2$\Delta$::URA3, SCS22$\Delta$::URA3, TCB1$\Delta$::hphNT1, TCB2$\Delta$::natNT2, IST2$\Delta$::kanMX6, TCB3-EGFP::HIS3MX6** | (S2B) | Lab stock |
| WKY0509 | *MAT$\alpha$, his3-$\Delta$200, leu2-3,112, ura3-52, lys2-801, TCB3$_{1-490}$-chimeraC-EGFP::HIS3MX6* | (3B-E,G-H), (S2E) | This study |
| WKY0527 | *MATa, his3-$\Delta$200, leu2-3,112, ura3-52, lys2-801, SCS2$\Delta$::URA3, SCS22$\Delta$::URA3, TCB1$\Delta$::hphNT1, TCB2$\Delta$::natNT2, IST2$\Delta$::kanMX6, TCB3$_{1-490}$-chimeraC -EGFP::HIS3MX6** | (3B-D,F-H), (S2E) | This study |
| WKY0529 | *MAT$\alpha$, his3-$\Delta$200, leu2-3,112, ura3-52, lys2-801, TCB3$_{191-490}$-chimeraC-EGFP::HIS3MX6* | (5B) | This study |
| WKY0530 | *MATa, his3-$\Delta$200, leu2-3,112, ura3-52, lys2-801, SCS2$\Delta$::URA3, SCS22$\Delta$::URA3, TCB1$\Delta$::hphNT1, TCB2$\Delta$::natNT2, IST2$\Delta$::kanMX6, TCB3$_{191-490}$-chimeraC-EGFP::HIS3MX6** | (5B,D-F,J), (S4A-C) | This study |
| WKY0531 | *MAT$\alpha$, his3-$\Delta$200, leu2-3,112, ura3-52, lys2-801, TCB3$_{191-1545}$-EGFP::HIS3MX6* | (S3A) | This study |
| WKY0562 | *MAT$\alpha$, his3-$\Delta$200, leu2-3,112, ura3-52, lys2-801, TCB3-EGFP::HIS3MX6, TCB1$\Delta$::URA3* | (1B) | This study |
| WKY0563 | *MAT$\alpha$, his3-$\Delta$200, leu2-3,112, ura3-52, lys2-801, TCB3-EGFP::HIS3MX6, TCB2$\Delta$::URA3* | (1B) | This study |
| WKY0587 | *MAT$\alpha$, his3-$\Delta$200, leu2-3,112, ura3-52, lys2-801, TCB2$\Delta$::URA3 TCB3$_{1-490}$-chimeraC-EGFP::HIS3MX6, TCB1$\Delta$:: hphNT1* | (3B-D) | This study |
| WKY0590 | *MAT$\alpha$, his3-$\Delta$200, leu2-3,112, ura3-52, lys2-801, TCB3-EGFP::HIS3MX6, TCB2-mRUBY2::kanMX* | (S1D) | This study |
| WKY0591 | *MAT$\alpha$, his3-$\Delta$200, leu2-3,112, ura3-52, lys2-801, TCB3$_{1-490}$-chimeraC-EGFP::HIS3MX6, TCB2-mRUBY2::kanMX* | (S2D) | This study |

| | | | |
|---|---|---|---|
| WKY0592 | *MATα, his3-Δ200, leu2-3,112, ura3-52, lys2-801, TCB3*$_{191-490}$*-chimeraC-EGFP::HIS3MX6, TCB2-mRUBY2::kanMX* | (S3C) | This study |
| WKY0595 | *MATα, his3-Δ200, leu2-3,112, ura3-52, lys2-801, TCB3-EGFP::HIS3MX6, TCB1-mRUBY2::kanMX* | (S1C) | This study |
| WKY0596 | *MATα, his3-Δ200, leu2-3,112, ura3-52, lys2-801, TCB3*$_{1-490}$*-chimeraC-EGFP::HIS3MX6, TCB1-mRUBY2::kanMX* | (S2D) | This study |
| WKY0597 | *MATα, his3-Δ200, leu2-3,112, ura3-52, lys2-801, TCB3*$_{191-490}$*-chimeraC-EGFP::HIS3MX6, TCB1-mRUBY2::kanMX* | (S3B) | This study |
| WKY0647 | *MATα, his3-Δ200, leu2-3,112, ura3-52, lys2-801, TCB3*$_{1-272}$*-chimeraC-EGFP::HIS3MX6, TCB2-mRUBY2::kanMX* | (S2F) | This study |
| WKY0648 | *MATα, his3-Δ200, leu2-3,112, ura3-52, lys2-801, TCB3*$_{1-272}$*-chimeraC-EGFP::HIS3MX6, TCB1-mRUBY2::kanMX* | (S2F) | This study |
| WKY0656 | *MATα, his3-Δ200, leu2-3,112, ura3-52, lys2-801, TCB3-ΔSMP-EGFP::HIS3MX6, TCB2-mRUBY2::kanMX* | (S1D) | This study |
| WKY0657 | *MATα, his3-Δ200, leu2-3,112, ura3-52, lys2-801, TCB3-ΔSMP-EGFP::HIS3MX6, TCB1-mRUBY2::kanMX* | (S1C) | This study |
| WKY0664 | *MATa, his3-Δ200, leu2-3,112, ura3-52, lys2-801, SCS2Δ::URA3, SCS22Δ::URA3, TCB1Δ::hphNT1, TCB2Δ::natNT2, IST2Δ::kanMX6, TCB3*$_{1-272}$*-chimeraC-EGFP::HIS3MX6** | (4A,C-E,G-H) | This study |
| WKY0665 | *MATa, his3-Δ200, leu2-3,112, ura3-52, lys2-801, SCS2Δ::URA3, SCS22Δ::URA3, TCB1Δ::hphNT1, TCB2Δ::natNT2, IST2Δ::kanMX6, TCB3*$_{191-272}$*-chimeraC-EGFP::HIS3MX6** | (5C-E,G-I) | This study |
| WKY0669 | *MATα, his3-Δ200, leu2-3,112, ura3-52, lys2-801, TCB3-ΔSMP-EGFP::HIS3MX6* | (1C,F,G,H) | This study |
| WKY0670 | *MATα, his3-Δ200, leu2-3,112, ura3-52, lys2-801, TCB1Δ::URA3, TCB2Δ::hphNT1, TCB3-ΔSMP-EGFP::HIS3MX6* | (S1F-H) | This study |
| WKY0672 | *MATa, his3-Δ200, leu2-3,112, ura3-52, lys2-801, SCS2Δ::URA3, SCS22Δ::URA3, TCB1Δ::hphNT1, TCB2Δ::natNT2, TCB3*$_{1-272}$ *-EGFP-chimeraC, IST2Δ::kanMX6* | (4B-D,F-H) | This study |
| WKY0675 | *MATα, his3-Δ200, leu2-3,112, ura3-52, lys2-801, TCB3-EGFP::HIS3MX6, TCB1-ΔSMP* | (1C), (S1E-G) | This study |
| WKY0676 | *MATα, his3-Δ200, leu2-3,112, ura3-52, lys2-801, TCB3-EGFP::HIS3MX6, TCB2-ΔSMP* | (1C) | This study |
| WKY0677 | *MATα, his3-Δ200, leu2-3,112, ura3-52, lys2-801, TCB3-EGFP::HIS3MX6 TCB1-ΔSMP, TCB2-ΔSMP* | (1C,F,G,I) | This study |

| | | | |
|---|---|---|---|
| WKY0678 | *MATα, his3-Δ200, leu2-3,112, ura3-52, lys2-801, TCB1-ΔSMP-EGFP::HIS3MX6* | (S1B) | This study |
| WKY0679 | *MATα, his3-Δ200, leu2-3,112, ura3-52, lys2-801, TCB2-ΔSMP-EGFP::HIS3MX6* | (S1B) | This study |
| WKY0680 | *MATα, his3-Δ200, leu2-3,112, ura3-52, lys2-801, TCB3$_{191\text{-}272}$-chimeraC-EGFP::HIS3MX6* | (5C) | This study |
| WKY0690 | *MATα, his3-Δ200, leu2-3,112, ura3-52, lys2-801, TCB3-ΔC2-EGFP::HIS3MX6* | (2A,C-E) | This study |
| WKY0691 | *MATα, his3-Δ200, leu2-3,112, ura3-52, lys2-801, TCB3-ΔC2-EGFP::HIS3MX6, TCB2-mRUBY2::kanMX* | (2B) | This study |
| WKY0692 | *MATα, his3-Δ200, leu2-3,112, ura3-52, lys2-801, TCB3-ΔC2-EGFP::HIS3MX6, TCB1-mRUBY2::kanMX* | (2B) | This study |
| WKY0693 | *MATa, his3-Δ200, leu2-3,112, ura3-52, lys2-801, TCB1-ΔC2-EGFP::HIS3MX6* | (2A) | This study |
| WKY0694 | *MATa, his3-Δ200, leu2-3,112, ura3-52, lys2-801, TCB2-ΔC2-EGFP::HIS3MX6* | (2A) | This study |
| WKY0716 | *MATα, his3-Δ200, leu2-3,112, ura3-52, lys2-801, TCB3-ΔC2-EGFP::HIS3MX6, TCB1-ΔSMP, TCB2-ΔSMP* | (2D,E,I) | This study |
| WKY0721 | *MATα, his3-Δ200, leu2-3,112, ura3-52, lys2-801, TCB3-EGFP::HIS3MX6, RTN1-mCHERRY::kanMX* | (2F) | This study |
| WKY0722 | *MATα, his3-Δ200, leu2-3,112, ura3-52, lys2-801, TCB3-ΔC2-EGFP::HIS3MX6, RTN1-mCHERRY::kanMX* | (2G) | This study |
| WKY0723 | *MATα, his3-Δ200, leu2-3,112, ura3-52, lys2-801, TCB3-ΔC2-EGFP::HIS3MX6, TCB1-ΔSMP, TCB2-ΔSMP, RTN1-mCHERRY::kanMX* | (2H) | This study |

* His marker is likely lost. All strains labelled with an asterisk are derived from WKY0270, which does not grow on -His plates. For WKY0527 and WKY0530, the loss of the His marker was further confirmed by PCR.

**Supplementary Table 2: Guide RNA forward sequences**

| Plasmid Number | CRISPR modifications | 20 nt gRNA sequence |
|---|---|---|
| pWK00319 | $Tcb3_{1\text{-}490}$-chimeraC-GFP<br>$Tcb3_{1\text{-}272}$-chimeraC-GFP<br>Tcb3-ΔC2 | AACATTCACCAACTCCTCAG |
| pWK0325 | $Tcb3_{191\text{-}1545}$ | GAGGAGAAGAAGAAAGTAGG |
| pWK0374 | $Tcb3_{1\text{-}272}$-chimeraC-GFP<br>$Tcb3_{1\text{-}272}$-GFP-chimeraC | CATATCCGAGACATCGCTTG |
| pWK0375 | Tcb3-ΔSMP | AACGTTAACCCTCAACTGGC |
| pWK0411 | Tcb1-ΔSMP | GATGTTGTTGTGATGGACTG |
| pWK0412 | Tcb2-ΔSMP | TCAAGATCAGACATGTCGTG |

**Supplementary Table 3: Templates for homology directed repair**

| **Mutation** | **Sequence** |
|---|---|
| Tcb3$_{1-490}$-chimeraC-GFP | ATATAGGGCCTATGCTATTCCCTCCGAACCATTTG<br>GATATTAATGTTGAAGACATTATGGCTGCTCAATTA<br>AAAGAAGCTATTGGTGGCGGAGGGGGCTCAGGG<br>GGAGGAGGTTCTGGGGGTGGAGGAAGTGGTGGG<br>GGTGGCTCAGGGGGAGGCGGATCCGCATCCAAG<br>GAGAAACCGAAACATAAAAAAGGCTTGCTGCATAA<br>GTTGAAAAAGAAACTTAAGGAAAAGCCTAAACACA<br>AGAAGGGTTTATTGCACAAATTGAAGAAAAAGCTG<br>CGTACGCTGCAGGTCGACGGAGCAGGTGCTGGT<br>GCTGGTGCTGGAGCAATGAGCAAGGGCGAGGAG<br>CTGTTCACCGGGGTGGTGCCCATCCTGGTCGAGC<br>TGGACGGCGACGTAAACGGCCACAAGTTCA |
| Tcb3$_{191-1545}$ | AATATAAAGAGCATCATTAGCTAGCTTTGTTTATTG<br>ACATACGCAATATTGCTAGAAAGAAAATAAGTAGA<br>GCTACGCATTGAAAGTCAAGAAAAATGTCCAGGGT<br>TATAAAAGCTTACATTCTGGAAAATTTTTATAACGA<br>TTGGTACTGTAATATAGCCACCGTTCTTGGAACTT<br>GTTTCTTCTCATGGTTATTTGCTTACATTGGGTTTT<br>CATGGTGGTCTATGATATTTATCTTCTTGGGAACT<br>GCGACCGTTTACAACGCAGAATATACAAGATTCAA<br>CAGAAATATCAGAGATGACTTGAAAAGAGTTACAG<br>TCGAAGAAACCTTGTCGGAT |
| Tcb3$_{1-272}$-chimeraC-GFP | GTAATATAGCCACCGTTCTTGGAACTTGTTTCTTCT<br>CATGGTTATTTGCTTACATTGGGTTTTCATGGTGG<br>TCTATGATATTTATCTTCTTGGGAACTGCGACCGT<br>TTACAACGCAGAATATACAAGATTCAACAGAAATA<br>TCAGAGATGACTTGAAAAGAGTTACAGTCGAAGAA<br>ACCTTGTCGGATCGCGGTGGCGGAGGGGGCTCA<br>GGGGGAGGAGGTTCTGGGGGTGGAGGAAGTGGT<br>GGGGGTGGCTCAGGGGGAGGCGGATCCGCATCC<br>AAGGAGAAACCGAAACATAAAAAAGGCTTGCTGC<br>ATAAGTTGAAAAAGAAACTTAAGGAAAAGCCTAAA<br>CACAAGAAGGGTTTATTGCACAAATTGAAGAAAAA<br>GCTGCGTACGCTGCAGGTCGACGGAGCAGGTGC<br>TGGTGCTGGTGCTGGAGCAATGAGCAAGGGCGA<br>GGAGCTGTTCACCGGGGTGGTGCCCATCCTGGTC<br>GAGCTGGACGGCGACGTAAACGGCCACAAGTTCA |
| Tcb3-ΔSMP | GTTACAGTCGAAGAAACCTTGTCGGATCGCTCAAA<br>AGAAGCTATTGGTGTCCTTGCCGTA |
| Tcb1-ΔSMP | TCATTATAAGTTTTCAATGGGGTCGGCCTTTTTTGT<br>CATCGTTATCACTTCATTGTTGTATAGAACCTCTGC<br>TAAAAAATACAGAGGTTCCATAAGAGAGTTAGTCC<br>AGAAAGAGTTCACGGTGCAGAAAGTGGAAAACGA<br>CTCTGGTTCCAACTTGTCCATTGGTATATTAGAAAT |

| | |
|---|---|
| | AACTGTCAAAAATGCAAAGGGGCTAAAACGTACCT CTTCGATATTGAACGAATCAATCGACCCTTACTTAT CTTTCGAATTTAACGATATATCTATTGCCAAGACAA GAACCGTAAGAGACACATTGAACCCC |
| Tcb2-ΔSMP | GCTGGGCTATTTTAAATTCTCTCTCGCGTCCGTCC TTATCGTGATGCTAACCACAGGAATGTTATATAGA ACTTCGTCCAAAAAATATAGGGAATCGTTAAGGGA TTTAGCCCAGAAAGAACAAACTGTAGAGAAAATTA CTAGTGATCTTTCCAAGACTGGCTTACCTATAGGT GTTTTGGAAATCAAAGTCAAAAATGCCCATGGATT AAGAAAACTTGTGGGCATGATCAAGAAAACAGTTG ACCCATACTTGACATTTGAGCTCTCTGGTAAAATA GTCGGTAAAACTAAGGTTTTTAAAAATTCTGCTAAT CCTGTTTGGAATGAATCC |
| Tcb3$_{1-272}$-GFP-chimeraC | AGAGATGACTTGAAAAGAGTTACAGTCGAAGAAAC CTTGTCGGATCGCCGTACGCTGCAGGTCGACGGA GCAGGTGCTGGTGCTGGTGCTGGAGCAATGAGCA AGGGCGAGGAGCTGTTCACCGGGGTGGTGCCCA TCCTGGTCGAGCTGGACGGCGACGTAAACGGCCA CAAGTTCAGCGTGTCCGGCGAGGGCGAGGGCGA TGCCACCTACGGCAAGCTGACCCTGAAGTTCATC TGCACCACCGGCAAGCTGCCCGTGCCCTGGCCC ACCCTCGTGACCACCCTGACCTACGGCGTGCAGT GCTTCAGCCGCTACCCCGACCACATGAAGCAGCA CGACTTCTTCAAGTCCGCCATGCCCGAAGGCTAC GTCCAGGAGCGCACCATCTTCTTCAAGGACGACG GCAACTACAAGACCCGCGCCGAGGTGAAGTTCGA GGGCGACACCCTGGTGAACCGCATCGAGCTGAA GGGCATCGACTTCAAGGAGGACGGCAACATCCTG GGGCACAAGCTGGAGTACAACTACAACAGCCACA ACGTCTATATCATGGCCGACAAGCAGAAGAACGG CATCAAGGTGAACTTCAAGATCCGCCACAACATCG AGGACGGCAGCGTGCAGCTCGCCGACCACTACC AGCAGAACACCCCCATCGGCGACGGCCccGTGCT GCTGCCCGACAACCACTACCTGAGCACCCAGTCC GCCCTGAGCAAAGACCCCAACGAGAAGCGCGATC ACATGGTCCTGCTGGAGTTCGTGACCGCCGCCGG GATCACTCTCGGCatGgACGAGCTGTACAAGGGTG GCGGAGGGGGCTCAGGGGGAGGAGGTTCTGGG GGTGGAGGAAGTGGTGGGGGTGGCTCAGGGGGA GGCGGATCCGCATCCAAGGAGAAACCGAAACATA AAAAAGGCTTGCTGCATAAGTTGAAAAAGAAACTT AAGGAAAAGCCTAAACACAAGAAGGGTTTATTGCA CAAATTGAAGAAAAAGCTGTAATAAACAGTGAAAA CTATTCTTTTATACGCTCAATAAGGGCAC |
| Tcb3-ΔC2 | AACCATTTGGATATTAATGTTGAAGACATTATGGCT GCTCAATCAAAAGAAGCTATTCGTACGCTGCAGGT CGACGGAGCAGGTGCTGGTGCTGGTGCTGGAGC |

| | |
|---|---|
| | AATGAGCAAGGGCGAGGAGCTGTTCACCGGGGT GGTGCCCATCCTGGTCGAGCTGGACGGCGACGT AAA |
